# Systematically characterizing the roles of E3-ligase family members in inflammatory responses with massively parallel Perturb-seq

**DOI:** 10.1101/2023.01.23.525198

**Authors:** Kathryn Geiger-Schuller, Basak Eraslan, Olena Kuksenko, Kushal K. Dey, Karthik A. Jagadeesh, Pratiksha I. Thakore, Ozge Karayel, Andrea R. Yung, Anugraha Rajagopalan, Ana M Meireles, Karren Dai Yang, Liat Amir-Zilberstein, Toni Delorey, Devan Phillips, Raktima Raychowdhury, Christine Moussion, Alkes L. Price, Nir Hacohen, John G. Doench, Caroline Uhler, Orit Rozenblatt-Rosen, Aviv Regev

## Abstract

E3 ligases regulate key processes, but many of their roles remain unknown. Using Perturb-seq, we interrogated the function of 1,130 E3 ligases, partners and substrates in the inflammatory response in primary dendritic cells (DCs). Dozens impacted the balance of DC1, DC2, migratory DC and macrophage states and a gradient of DC maturation. Family members grouped into co-functional modules that were enriched for physical interactions and impacted specific programs through substrate transcription factors. E3s and their adaptors co-regulated the same processes, but partnered with different substrate recognition adaptors to impact distinct aspects of the DC life cycle. Genetic interactions were more prevalent within than between modules, and a deep learning model, comβVAE, predicts the outcome of new combinations by leveraging modularity. The E3 regulatory network was associated with heritable variation and aberrant gene expression in immune cells in human inflammatory diseases. Our study provides a general approach to dissect gene function.

## INTRODUCTION

Despite systematic efforts in genetics and genomics, our knowledge of the function of many genes remains limited, especially those from large gene families, where the general molecular function may be inferred from sequence features, but the specific mechanism, biological process, cellular context and physiological impact of individual genes and their combinations often remain partly or completely unknown. Multiple approaches can help decipher individual gene function, including Genome-Wide Association Studies (GWAS) to relate causal genetic variants to quantitative traits^1^; forward genetic screens followed by phenotypic assessment, including cell viability, images or molecular profiles^2^; and guilt-by-association approaches, based on similarity in molecular patterns between a gene of interest and other genes. Despite their power and utility, each of these approaches has some limitations. Genetic association studies are often limited by the modest effect sizes associated with common variants in human populations^3^; correlative approaches provide suggestive associations but not causal relations; and forward genetic screens have typically had to pre-define the phenotype of interest, such as cell viability^4^ or a cellular marker^5, 6^. Finally, all approaches are challenged at deciphering genetic interactions, due to limited statistical power or experimental scale to test exponentially large numbers of combinations.

Large protein families, such as E3 ubiquitin ligases (“E3s”), are an important example of this challenge. The human genome codes for >600 different E3s responsible for catalyzing the ligation of ubiquitin (Ub) to substrates in almost every biochemical pathway^7^, including many immune functions^8^. GWAS have implicated variants in E3 ligase genes in many diseases, including inflammatory and autoimmune diseases^9, 10^, but characterizing their specific cellular roles remains challenging, as is determining their inter-relationships. In particular, dendritic cells (DCs) play a key role in initiating immune responses, including multiple inflammatory and autoimmune diseases^11, 12^, and heritable variants in multiple genes in DCs contribute to their aberrant inflammatory signaling in disease, including several axes that may be targeted therapeutically^13^. While previous studies implicate different E3 ligases in the DC inflammatory response to lipopolysaccharide (LPS)^5^, relatively little is known about the E3 circuit in these or other primary immune cells, as many studies focus on transformed cancer cell lines.

Recent advances in combining pooled genetic perturbation screens with rich, single cell readouts, especially single cell RNA-seq (scRNA-seq) in Perturb-Seq assays, open opportunities to dissect the function of genes and gene combinations from large gene families^14–24^. In Perturb-Seq screens, the perturbed genes can be partitioned into co-functional modules, based on the similarity of their effects across many genes, controlling co-regulated programs of genes responding similarly across multiple perturbations^16, 17^. Moreover, any diversity in cell subsets or processes, such as the cell cycle or differentiation, is naturally captured in the screen^16^. Most Perturb-Seq studies to date, and especially the very few done in primary cells, have analyzed up to a few dozen perturbations^14–22^, with a recent notable screen of thousands of genes but only in a transformed cell line^23^.

Here, we used Perturb-seq at scale to screen the function of each of 1,130 genes spanning E3 ligases, E3-like proteins and their interacting partners and substrates in the inflammatory response to stimulation with LPS in primary mouse bone marrow derived dendritic cells (BMDCs). The cells in one experiment spanned DC1, DC2, and migratory DC (mDC) subsets, a gradient of DC maturation, and a range of gene programs, allowing us to decipher the role of E3 ligases in multiple contexts simultaneously. A regulatory model distinguished six co-functional modules impacting eleven different programs of co-regulated genes, showing which E3 ligases, adaptors and substrate recognition adaptor proteins regulate each process in DCs, capturing known associations and making many new functional annotations. Computational integration of the regulatory (genetic) model with physical protein-protein interactions and transcription factor (TF)-target genes relations shows that the regulatory model was congruent with physical mechanistic interactions. E3s and their physically interacting partners were enriched in the same co-functional module, and the programs they regulated were enriched for targets of specific TFs, highlighting multiple paths from E3s and complex members through TFs to different DC processes they regulate. Moreover, the circuit was modular, such that Cullin-RING ligases (the largest subfamily; >200 members) and their known adaptors co-regulated the same processes, but combined with different substrate recognition adaptor proteins to control distinct aspects of the DC life cycle. The circuit was also congruent with human disease biology, with both co-functional modules and their regulated programs enriched in heritability for risk immune and inflammatory diseases. Leveraging our large screen’s design to also randomly sample combinations of perturbations, we found that intra-module (non-additive) genetic interactions are more prevalent than inter-module ones and then used the modular architecture to devise com VAE, a new deep learning model that predicts genetic interactions. Our study offers a general scalable approach to dissect gene function, including physiological functions for dozens of E3s and related genes, congruent physical circuits, principles of modularity in the regulatory and molecular architecture, characterization and prediction of genetic interactions, and an overall model of the inflammatory response to help interpret human genetics signal at unprecedented resolution.

## RESULTS

### A systematic screen of E3 ligases in immune dendritic cells

To study the role and circuitry of members of the large gene family of E3 ligases in inflammatory responses, we curated a comprehensive set of 1,137 genes encoding E3s and related proteins to screen them by Perturb-seq (**Table S1-2**; **STAR Methods**). These included 382 genes with ‘E3 activity’ designation in the Integrated Annotations for Ubiquitin and Ubiquitin-like Conjugation Database (iUUCD ^7^), such as proteins from RING, HECT, U-box, PHD, RBR, and other families; 509 genes with ‘E3 adaptor’ designation in iUUCD, such as those from DWD, BTB, APC, Cullin, BC-box, F-box, DDB1, and other families; 6 genes with an annotated ubiquitin binding domain and one with a ubiquitin-like domain (Rbbp6) also from iUUCD; and 239 genes based on an NCBI search for ‘E3 activity’, capturing other enzymes in the ubiquitylation cascade (E1s, E2s), known E3 substrates (*e.g.*, Tp53, Ikbke, and Cebpb), and members of relevant signaling networks regulated by E3 ligases (*e.g.*, TLR and TNF signaling). We synthesized and screened guides targeting 1,130 of the genes (we could not confidently design gRNAs for 7 putative pseudogenes).

We optimized Perturb-seq for large scale screening and used it to screen the 1,130 genes, perturbed by 3,390 targeting guides, profiling 838,201 individual BMDCs, after 3 hours of treatment with LPS (**Figure 1A**). We chose the 3h time point because the DC transcriptional response to LPS has a single wave, peaking around 3hr for mature mRNA at both the population and single cell level, and is optimal for observing RNA expression effects^5, 25–30^, whereas protein level changes occur later^26^. We isolated 54 million cells from the bone marrow of Cas9 transgenic mice^31^, treated them with GM-CSF to differentiate them towards BMDCs and, on day 2, transduced them at a planned multiplicity of infection (MOI) of 0.2 with a pooled lentiviral library of 3,390 sgRNAs targeting the 1,130 genes (3 guides per gene) and 330 control guides (165 targeting intergenic regions and 165 non-targeting). We designed a new Perturb-Seq vector (pRDA122; **Figure 1A**) with a capture sequence appended to the 3’ terminus of the gRNA scaffold sequence to enable direct capture of CRISPR gRNAs for scRNA-seq (**STAR Methods**), compatible with feature barcoding for droplet-based 3’ scRNA-seq^24^. We continued to differentiate the transduced cells for another 7 days, when they are predominantly BMDCs^25, 30, 32, 33^, and then treated them with LPS. At 3 hours post-LPS treatment, we sorted mKate2^+^Cas9-2A-EGFP^+^ cells, loaded 2.32 million onto 46 channels using cell hashing and “super-loading”^34, 35^ (**Figure S1A-D**), and performed scRNA-seq. We obtained profiles from 1,071,671 non-empty droplets (EmptyDrops ^36^ FDR < 0.01, **STAR Methods**), containing 838,201 single cells and 233,470 multiplets, followed by dedicated PCR to detect the guide RNA in each cell (**STAR Methods**). After QC and guide assignment, we retained 519,535 single cell profiles assigned with one or more gRNA (targeting or control; detected MOI of 1.2) for our main analyses, followed by analysis of 177,871 cells with multiple perturbations with dedicated methods below, as well as applying a compressed sensing approach to all the multiplets in a companion study (Yao et al, bioRxiv 2023). As reference, we also profiled a total of 10,347 unperturbed cells, encompassing both LPS stimulated and unstimulated cells.

**Figure. 1.**
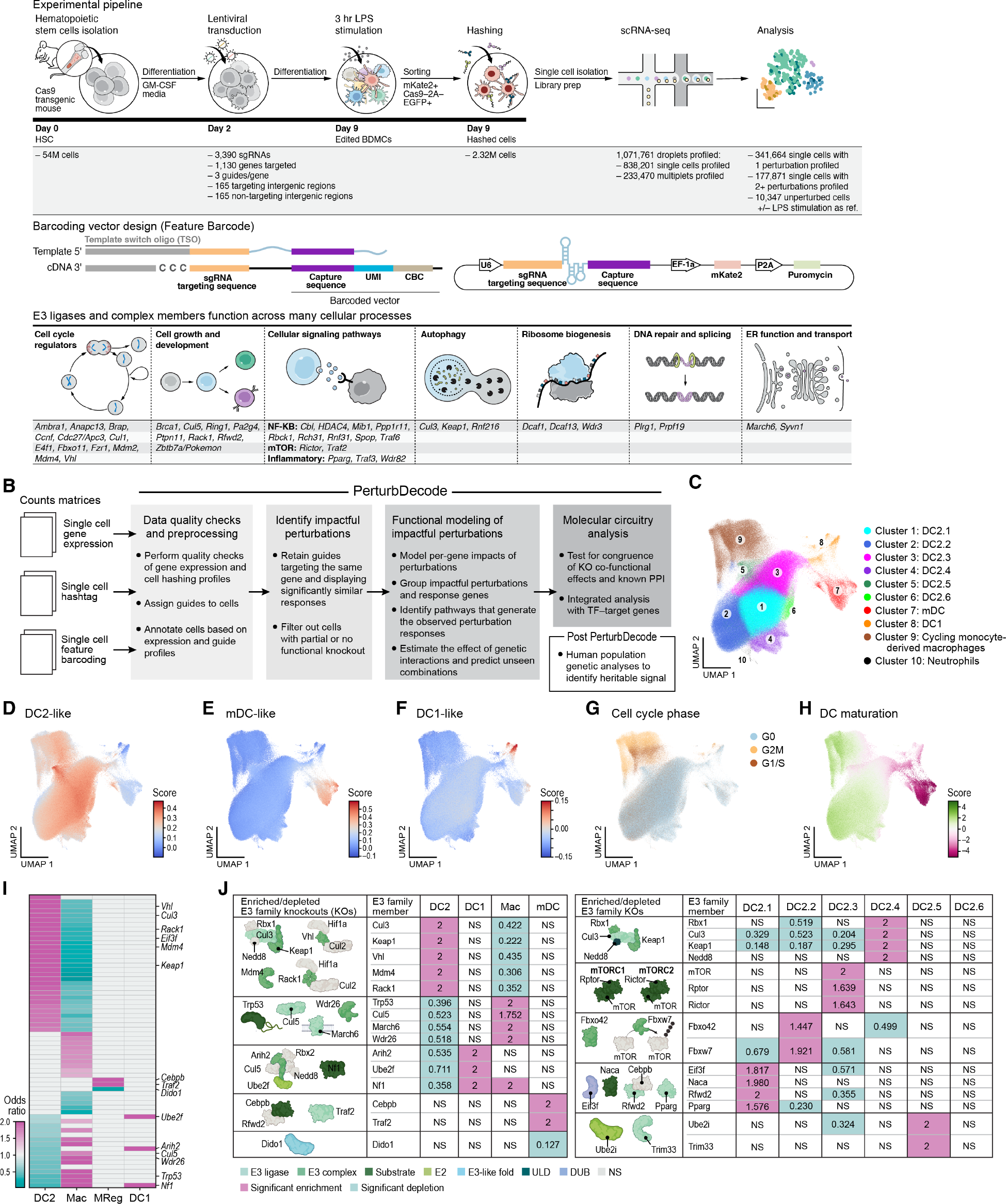
Large scale Perturb-seq of the function of E3 ligases in the LPS response in BMDCs **A.** Study overview. Top: Experimental flow. Middle: Perturb-Seq vector design. Bottom: Example E3 ligases and complex members known to regulate different processes. **B.** PerturbDecode. Workflow diagram of key features (full diagram in **Figure S1E**). **C-E.** Screened cell populations captures multiple cell states and subtypes. Uniform Manifold Approximation and Projection (UMAP) of 519,535 cell profiles colored by cluster membership (**C**), cell type signature scores (**D-F**), cell cycle phase (**G**), or the difference of macrophage vs. DC signature scores (**STAR Methods**) (**H**). **I-J**. E3 perturbations affect DC subset distributions. **I.** Odd-ratio (color bar) of the significant (FDR < 0.15, one-sided Fisher’s exact test) enrichment (pink) or depletion (blue) of guides targeting perturbed genes (rows) in each major cell subset (columns) in the screen. **J.** Summary of enrichment (pink) and depletion (blue) of guides targeting key proteins in major subsets (left) and in DC2 subtypes (right), colored by E3 family type and grouped by complex. ULD: ubiquitin-like domain; DUB: de-ubiquitylating enzyme; NS: non-significant. See also related Figures S1 and S2

### An end-to-end computational pipeline for large Perturb-seq screens

To analyze large screens, we developed PerturbDecode, for end-to-end, automated analysis in four consecutive pillars (**Figure 1B and S1E, STAR Methods**): (**1**) data QC and preprocessing; (**2**) identification of the effects of perturbations on genes; (**3**) learning the regulatory topology of perturbed and impacted genes for single and/or combinatorial perturbations; and (**4**) relating the regulatory (genetic) topology to physical interactions and human genetics (**STAR Methods**). Briefly, in pre-processing, PerturbDecode addresses hashing and feature barcoding assignment, detects depleted feature barcodes and cells with multiple guides and removes outliers. Next, it efficiently identifies impactful guides and impacted cells^16^, by estimating the effect of each guide on each gene with a negative binomial linear regression model, accounting for confounders (**STAR Methods**). It retains impactful guides defined as those with effects that are significantly similar to those of at least one other guide targeting the same gene (*vs*. a background of all guides), and iteratively identifies and retains impacted cells^16^. To determine the impact at the level of perturbed genes, PerturbDecode uses a mixed effects negative binomial linear regression model, with cell subsets inferred by initial clustering as random effects and the feature barcode matrix as the set of fixed effects, correcting for confounders (**STAR Methods**). It retains perturbations affecting a significant (FDR < 0.1) number of genes (**STAR Methods**), clusters the resulting coefficient matrix to generate co-functional modules and co-regulated programs, and decomposes the matrix by independent component analysis (ICA) to infer latent independent processes that could generate the observed perturbation responses. For combinatorial perturbations, it estimates the effect of genetic interactions and predicts the impact of unseen combinations. Finally, it includes multiple *post-hoc* analytics to relate the learned model to molecular circuits (protein-protein and TF-target interactions) and to human genetics data.

### Perturb-Seq screen yields impactful perturbations consistently across guides

Estimating the impact of each of the 3,204 targeting guides (detected in at least 20 cells) on the expression of each of 6,685 genes (expressed in at least 5% of cells) showed that the correlation in effect sizes between cells with guides targeting the same gene was significantly higher than between guides targeting different genes or between targeting and control guides (P-value < 10^-16^, Kolmogorov-Smirnov (KS) test, **Figure S1F**). Focusing on concordant guides (**STAR Methods**), we retained 2,263 KO guides targeting 1,031 genes for downstream analysis and learned a model at the targeted gene (rather than guide) level (**STAR Methods**). The average number of genes significantly affected by each perturbed gene (FDR < 0.1, **STAR Methods**) was significantly higher than for controls (36.91 *vs*. 0.78 (non-targeting) and 1.04 (intergenic) on average, P < 2.2*10^-16^, one-sided Wilcoxon rank-sum test, **Figure S1G, STAR Methods**). Of the 1,031 targeted genes, 544 were also among the 6,685 analyzed genes: the vast majority had 491 had a nominal negative effect on their own expression, 137 of them significantly (FDR< 0.1, **Figure S1H**), and only four had a significant positive effect. Expressed (detected) E3s affected significantly more genes than undetected ones (P < 10^-4^, one-sided non-parametric Wilcoxon test, **Figure S1J**) and the mean expression of the targeted (KO) gene in unperturbed cells was modestly but positively correlated with the number of genes impacted by their perturbation (Spearman’s ρ = 0.22, P < 1.5*10^-10^, **Figure S1I**). Overall these results suggest that CRISPR-induced indels overall caused nonsense mediated decay (NMD) of the respective transcripts, for expressed E3s and family members, as well as with prior observations that some CRISPR-knockout generated indels having poor NMD ^5, 16^ or even (futile) transcriptional compensation^37^.

### DC1-, DC2-, migratory DC-, and macrophage-like cells are screened jointly

BMDC populations are heterogeneous, and previous studies^29, 30, 38–40^, including our earlier Perturb-Seq screen in this system^16^, all highlighted the presence of different subsets, including cells expressing macrophage-like signatures, and “cluster-disrupted” DCs^30, 41^. Because Perturb-Seq characterizes any cell diversity *post hoc*, it assesses these multiple phenotypes simultaneously^16^.

The 519,535 perturbed single cell profiles partitioned into ten clusters (**Figure 1C-H** and **S2A**, **Table S3** and **S4**), which included multiple DC2-like subsets (high expression of Cd9, Il1b, and Sirpa (but also Irf4 and Il6); **Figure 1C,D,** clusters 1-6; **Figure S2A,B**); migratory DC (mDC)-like cells (high expression of Ccr7, Fscn1, Il4i, Socs2, and Relb (but not Pdl2); cluster 7, **Figure 1C,E** and **S2A,C**); DC1-like cells (high expression of Clec9a, Xcr1, Batf3, Irf8, Tap1, Flt3, and Wdfy4 but also Cd8a and Tlr3, and no expression of pDC marker genes (Tcf4, Tlr7, and Tlr9; additionally, Siglec-H was not among detected transcripts) **Figure 1C,F** and **S2A,D,E**), and macrophage-like cells (high expression of M1 markers Il6, Il1b, and Fpr2 and moderate expression of M2 markers Chil3, Fn1, and Mc1; **Figure 1C** and **S2A,F,G**). The three main DC subsets also followed a gradient of expression from more mature DC-like (most prominent in mDCs) to macrophage-like (more prominent in some DC2s) signatures (**Figure 1H** and **S2H,I**). Each DC2-like cell subset had additional distinguishing markers (**Figure S2A**). Cycling cells (**Figure 1C,G**, Cluster 9, 13%) expressed signatures of either DC2s or macrophages/monocytes (**Figure 1D,G,H** and **S2H,I**). These cycling monocyte-derived macrophages are consistent with previous reports^38, 42^. A very small subset (460 cells; 0.08%) expressed a neutrophil signature (**Figure 1C** and **S2A**, cluster 10). DC2-like, DC1-like, mDC-like, and cycling monocyte-derived macrophages were also present in genetically unperturbed cells, with and without LPS stimulation (**Figure S2J-W**), but at different proportions, with the fraction of macrophage-like cycling cells lower in genetically unperturbed LPS stimulated cells than in either unperturbed unstimulated DCs or perturbed stimulated DCs (**Figure S2W**, P-value < 2.2*10^-16^, one-sided Fisher’s exact test, **STAR Methods**), and the fraction of mDCs in unperturbed cells (stimulated and unstimulated) higher than in perturbed stimulated DCs (**Figure S2W**, P-value < 2.2*10^-16^, one-sided Fisher’s exact test). The increase in monocyte-derived macrophages in genetically perturbed cells (**Figure S2W**), suggests that the macrophage-like state is typically repressed by LPS stimulation but remains accessible upon perturbation.

### Multiple E3 ligases impact specific DC cell subsets or differentiation

Sixty-five (65) genes were targeted by two or more guides that were significantly depleted from BMDCs *vs*. the input guide library distribution, suggesting that these genes are essential for BMDC survival and proliferation^5, 16^ (**Table S5**, **STAR Methods**; P-value<0.05, considering a background of the corresponding change in control guides). Indeed, these were enriched for regulation of cell division (*e.g.*, Aurka, Myc, Plk1, Pou5f1, Prc1, Tle6, Wdr5, Ybx1), including Mdm2 (all three guides depleted), an E3 ligase that ubiquitylates p53 as an active heterodimer with Mdm4 and is essential for cell cycle regulation^43^. Some perturbations affected the proportions of all cycling cells of a specific type (**Figure S2X**), such as enrichment in cycling macrophages of guides targeting the tumor suppressor and E3 ligase substrate Trp53, and enrichment in cycling DC1-like cells of guides targeting the substrate Nf1, a tumor suppressor in myeloid cells that affects growth sensitivity to GM-CSF^44^ and regulates proliferation regulators^45^. Because different subsets were enriched for cycling cells (**Figure 1G**), some of these perturbations also shifted subset proportions. For example, consistent with previous reports^46^, Trp53 KO were enriched in cycling monocyte-derived macrophages, and depleted in non-cycling DC2s; and its negative regulator E3 ligase Mdm4 had the opposite pattern (**Figure 1J** and **S2X**).

Perturbation in 64 genes, including 29 E3s and complex members significantly affected the relative proportion of the main cell subsets, especially the balance of DC2s *vs*. cycling macrophage-like cells (**Figure 1I,J**, **STAR Methods, Table S6**; FDR < 0.15, one-sided Fisher’s exact test). Guides depleted in cycling macrophage-like cells *vs*. DC2s included the cullin E3 ligase Cul3 and its substrate adaptor Keap1, which ubiquitylates Nrf2, regulating redox response and inflammation^47, 48^; Vhl and Rack1, which togerher complex with Cul2^49^ and ubiquitylate Hif1a; and Mdm4, which interacts with Mdm2 to stimulate p53 ubiquitylation^50^. Vhl, Cul3, Keap1 and Mdm4 were not previously described as regulators of DC2 differentiation, but deletion of Nrf2 impacts both tissue resident and BMDC functions^51^ and Rack1 depletion in myeloid cells protects against viral infection in mice without affecting the number of F4/80^+^CD11b^+^ myeloid cells^52^. Conversely, guides enriched in cycling monocyte-derived macrophages *vs*. DC2s included those targeting Cul5, March6, and Wdr26, E3s not previously recognized as regulators of DC differentiation (**Figure 1I,J**).

Other enrichments and depletions further highlight the roles for E3s and related proteins in specific cell subsets. For example, mDCs were enriched for guides targeting the E3 ligase Traf2 and the TF CEBPB and depleted for guides targeting DIDO. Traf2 plays a role in TNF-mediated NF-KB and MAP kinase signaling, and TNF injection can increase DC trafficking to lymph nodes^53^, suggesting a potential role for Traf2 in regulating DC migration. Enrichment of guides targeting CEBPB in mDCs is consistent with our earlier findings and CEBPB’s natively low level in those cells (**Figure 1I,J**)^16^. Concomitantly, DC2.2s where CEBPB expression is higher, are enriched for guides targeting Rfwd2 (COP1), which in turn targets CEBPB for degradation^54^. Conversely, guides targeting Dido1, a gene previously characterized only in embryonic stem cell differentiation^55, 56^, were depleted in mDCs, suggesting a novel role in mDC differentiation (**Figure 1I,J**). In another example, guides targeting the E3 ligase Arih2, the E3 ligase substrate Nf1, and the E2 enzyme Ube2f were enriched in DC1s and macrophages and depleted in DC2s (**Figure 1I,J**). This is consistent with and expands on their established roles: Arih2 ubiquitylates substrates of Ube2f, the Nedd8 E2 that mediates neddylation of Cul5-Rbx2^57^, causing degradation of the NF-KB inhibitor Ikbb in DCs^58^, and inactivating mutations in Nf1 lead to uncontrolled cell growth and plasmocytoid DC (pDC) neoplasms^59^.

Many guides, including those targeting members of the WD-repeat protein subfamily, were specifically enriched in different DC2 subsets, including multiple members of a single complex in the same subset, suggesting that distinct DC2 subsets are controlled by different pathways (FDR < 0.15, one-sided Fisher’s exact test, **Figure 1J** and **S2Y**, **Table S6**). For example, perturbations of each of the four members of the KEAP1:NEDD8-CUL3:RBX1 complex were specifically enriched in DC2.4s. DC2.3s were enriched for guides targeting E3 ligase substrates and members of the mTOR pathway, including mTORC2 (Mtor and Rictor) and mTORC1 (Mtor and Rptor), consistent with and refining the role of mTor signaling in regulating differentiation and immune functions in DCs^60^. DC2.2s were enriched for guides targeting Fbox E3 ligase component members, including Fbxo42, a regulator of the TAK1 pathway that activates p38^61^, and Fbxw7, a regulator of antiviral immunity in macrophages^62^. DC2.1s were enriched for guides targeting Eif3f, Naca, Rfwd2 (Cop1), and Pparg. In addition to its role as a TF, Pparg is an E3 ligase that targets p65^63^ and regulates lipid metabolism in DCs, leading to ferroptosis and impairing maturation^64^, and DC-T cell interactions in type-2 immunity^65^. DC2.5s were enriched for guides targeting Trim33, an E3 that interacts with Pu.1 (SPI1) and regulates the NLRP3 inflammasome, LPS response, and macrophage activation^66, 67^, as well as for guides targeting Ube2i (Ubc9), an E2 SUMO-conjugating enzyme with a role in lupus, regulating sumoylation-based suppression of type I IFN and pDCs^68^. Thus, we identified regulators of six DC2 substates, including multiple complex members similarly associated with the same state(s).

### Six co-functional modules of E3 ligases regulate eleven gene programs

To relate these broad changes to regulatory mechanisms, we next learned a regulatory model associating 329 impactful perturbed genes (affecting the level of at least 15 genes) to 1,041 significantly impacted targets (affected by at least four of the 329 perturbations), and clustered the perturbed genes and impacted targets (**STAR Methods**) into six co-functional gene modules (M1-6) and eleven co-regulated gene programs (GP1-11), respectively (**Figure 2** and **Table S7-8, STAR Methods**).

**Figure 2.**
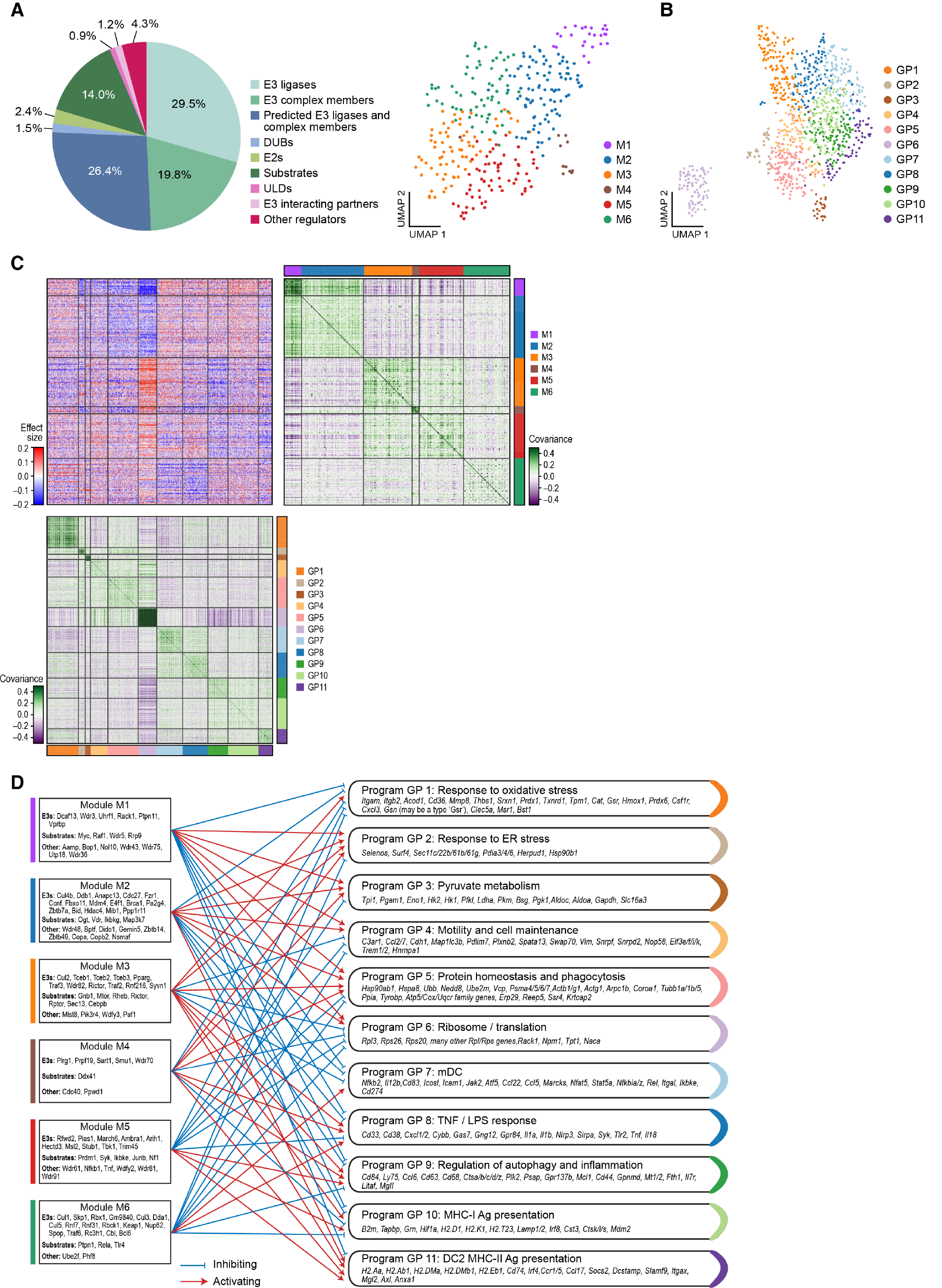
Six co-functional modules of E3 and related genes regulate 11 gene programs in the response of BMDCs to LPS **A.** Co-functional modules. Left: Distribution of E3 family member types across the 329 significant regulators. Right: UMAP embeddings of the regulatory profiles of the 329 regulators (KO genes), colored by their module membership (**STAR Methods**). **B.** Co-regulated programs. UMAP embeddings of the regulated profiles of 1,041 affected genes, colored by their gene program membership (**STAR Methods**). **C,D.** Regulatory model. **C.** Top left: Regulatory matrix (beta). Regulatory effect size (red/blue) of perturbing (KO) each of 329 genes (rows) on the expression of each of 1,041 affected genes (columns). Red/blue: Induction/repression in response to perturbation (KO) compared to control cells. Black horizonal and vertical line delineate co-functional modules and co-regulated programs, respectively. Top right: Co-functional modules. Covariance (green/purple) between the regulatory profiles in beta of perturbing each of 329 genes. Genes are clustered by module (as in **A**; color code on top and right). Bottom left: Co-regulated programs. Covariance (green/purple) between the regulatory profiles in beta of the effect on expression of each of 1,041 genes. Genes are clustered by program (as in **B**; color code on bottom and right). **D.** Regulatory network. Bipartite graph from co-functional modules to the co-regulated programs. Red point arrow/blue blunt arrow: module genes activate/inhibit program (*i.e.*, KO inhibits/activates program) (arrow color determined by significant mean difference). Key gene names are noted. See also related **Figure S3**.

The eleven programs (**Figure 2C,D** and **S3A-K, Table S8**) consisted of genes strongly co-regulated across the perturbations and were enriched for different immune and cellular processes. We annotated the programs by their enrichment in functional categories and previously described signatures (**Table S3,8**) as response to oxidative stress (GP1), ER stress response (GP2), pyruvate metabolism (GP3), motility and cell maintenance (GP4), protein homeostasis and phagocytosis (GP5), translation (high in mDCs and DC1s) (GP6), mDCs (GP7), TNF/LPS response (GP8), autophagy regulation and inflammation (GP9), MHC-I antigen presentation (GP10), and DC2 and MHC-II antigen presentation (GP11)^39^. Some programs are expressed broadly across all cell subsets (*e.g.*, GP5, **Figure S3E,M**) and others are quite specific (*e.g.*, GP6 in DC1s and mDCs; **Figure S3F,M**). The programs were not necessarily independent of each other: some were regulated in anti-correlated ways (*e.g.*, opposite regulatory effects on GP6 (DC1/mDC expressed translation) *vs*. GP9 (autophagy and inflammation) and 11 (DC2 and MHC-II presentation)), while others were regulated in similar (though not identical) ways (*e.g.*, GP4 (motility and cell maintenance), GP5 (protein homeostasis and phagocytosis), and GP6 (translation)) (**Figure 2C**).

The programs refined ones we previously defined in the same system under perturbation of 24 TFs^16^ (*e.g.*, P2 genes partition into mDC (GP7) and translation (GP6)) and uncovered new functional groupings (*e.g.*, DC2 MHC-II antigen presentation (GP11), response to ER stress (GP2), and pyruvate metabolism (GP3)) (**Figure S3L**), showing that the expanded scope and nature of perturbations could reveal additional regulatory processes.

### Regulatory patterns and targets reveal the functional roles and co-associations of E3s

The six co-functional modules of 329 regulators each impacted a different subset of programs (**Figure 2C,D**), such that modules M1 and 2 were generally positively correlated with each other and negatively correlated with modules M3, 4, and 5, reflecting opposing functions.

The co-functional modules included 97 E3 ligases, 65 E3 complex members and 3 ubiquitin-like domain proteins, 85 of which did not have prior literature evidence in DCs or inflammation, and can be deemed as novel functional annotations (**Table S9**), based on their co-membership and their target programs (**Figure 2D**). Three of the E3 ligase or complex members in the model are “authorities” in the E3 circuit, as their expression is impacted (positively and negatively) by a particularly high number of other E3s (**Figure 3A** and **S4A,B**): E3s Rack1 (which targets Hif1a and BimEL^49, 52, 69^) and Rnf128 (Grail; regulates Tbk1 and interferon and antiviral response^70–72^), and E3 complex member Socs3 (regulates Jak2 and gp130 and response to cytokines). All three have known roles in DC biology or inflammation: Rack1 suppresses type 1 IFN responses rendering mice more susceptible to viral infection^52^, Rnf128 targets the DC maturation marker Cd83 for degradation^72^, and Socs3 regulates JAK/STAT signaling, inflammation and macrophage polarization^73^. Notably, only 14 of 329 regulators were previously identified in a genome-wide CRISPR screen for regulators of TNF protein expression in the same system^5^, and these were further distinguished by our model into different modules, with most (9 of 14) in M6 (*e.g.*, Ube2f, Rbck1, Rnf31, Spop, Traf6, and Nedd8 (positive regulators), and Rc3h1 (negative regulator)). This shows the expanded power of functional discovery based on comprehensive expression profiles compared to one highly validated reporter/marker.

**Figure 3.**
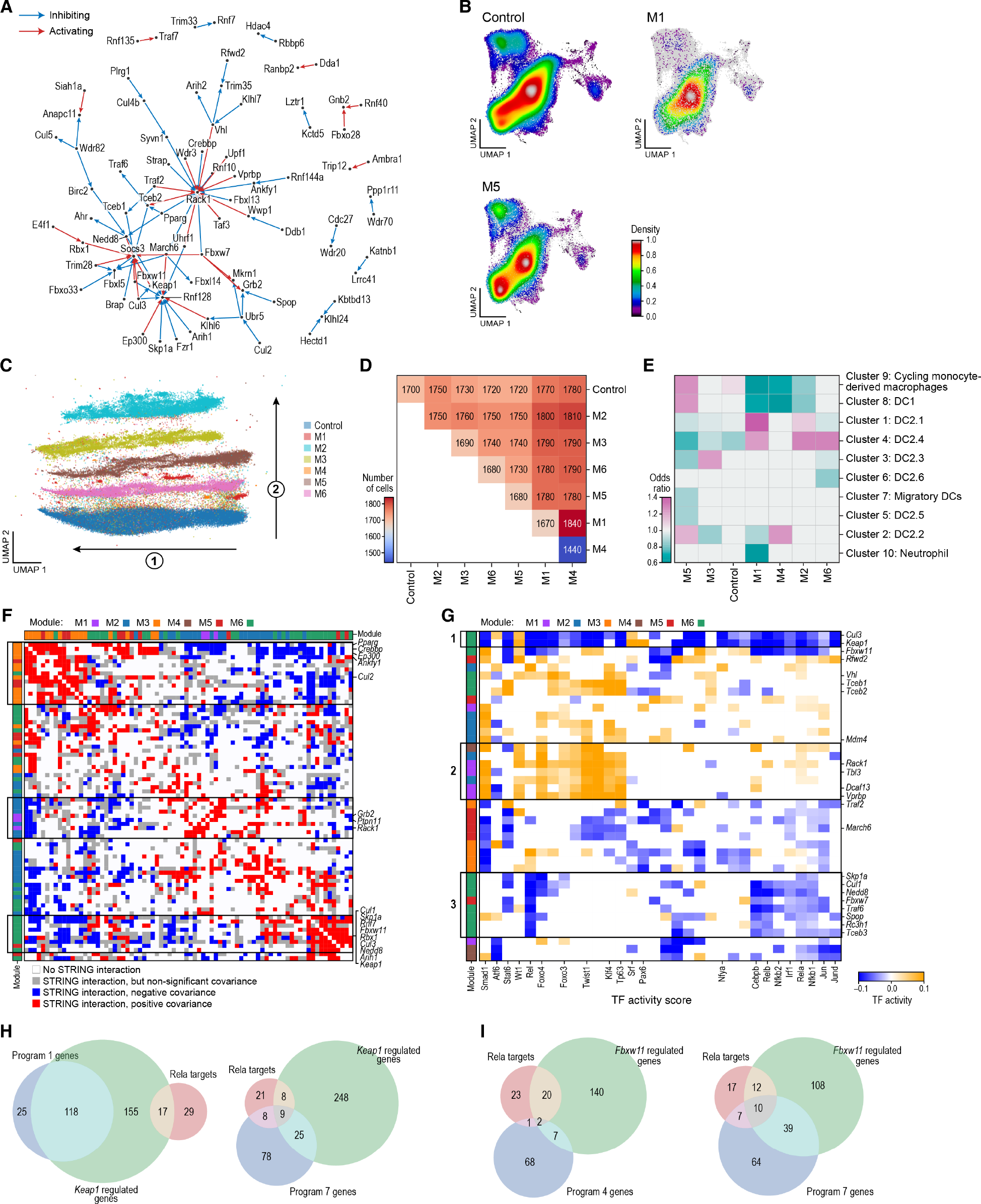
The genetic regulatory network is congruent with physical interactions and highlights modularity in E3-based regulation **A.** Three E3 ligases are highly regulated ‘authorities’ in the E3 network. Regulatory relations from the model (Figure 2C) based on a perturbation (KO) in an E3 ligase to another E3 ligase whose expression is affected. Red/blue arrows: perturbed E3 activates/inhibits target’s expression (*i.e.*, KO inhibits/activates expression). **B-E.** Co-functional modules impact distribution of cell states. **B.** UMAP embedding of single cell profiles colored by Gaussian kernel density estimations of cells control (top left), M1 (top right) or M5 (bottom) guides. **C.** Supervised UMAP embedding of DC2.1, DC2.2, and DC2.3 cell profiles using a cells’ co-functional module assignment as the response label (**STAR Methods**), colored by the module assignment of their guides (color code). **D.** Average Wasserstein distances (color bar) between cells with guides from different co-functional modules (rows, columns). **E.** Odd-ratio (colorbar) of significantly enrichment (pink) or depletion (green) of DC subsets (rows) in cells with guides from each co-functional module (columns) (FDR < 0.15, one-sided Fisher’s exact test). **F.** Co-functional modules are enriched for protein-protein interaction partners. Physical interactions (blue, red, grey; experimental score > 0, STRING DB) or lack thereof (white) between each pair of 78 E3 ligases and adaptors with at least 24 interactions (with any of the 165 E3 ligases among the 329 regulators). Red/blue: the regulatory profiles of the physically interacting genes have significant (P<0.05) positive/negative correlation. Grey: the regulatory profiles of the physically interacting genes are not significantly correlated. Bars: co-functional modules. Rows and columns are hierarchically clustered. Full matrix in **Figure S4G**. **G,H,I.** TF explaining expression impact of E3 perturbations. **G.** Inferred activity scores (color bar) of 32 TFs (columns) whose target genes are significantly (FDR < 0.1) induced (yellow) or repressed (blue) when perturbing each of 41 E3 and related genes (rows) (**STAR Methods**). (Full matrix in **Figure S4J**). **H,I.** Intersection between TF targets (DoRothEA), E3 expression targets, and gene programs. See also related **Figure S4**.

M1 regulators strongly activated ER stress (GP2), protein homeostasis and phagocytosis (GP5) and translation (GP6) through the action of regulators of ribosome biogenesis (predicted E3s Bop1, Wdr36 and Nol10, Cul4 substrate receptor Dcaf11 (VprBP); processome members E3 adaptors Dcaf13 and Wdr33), including those involved in the stress response (predicted E3s Wdr43, Wdr75, and Utp18), and cellular migration (E3 Aamp, predicted E3 Bop1) (**Table S9**). This is consistent with the enrichment of ribosome biogenesis and stress genes in the translation (GP6) and ER stress (GP2) programs. Additionally, M1 regulators repressed the mDC (GP7), TNF/LPS response (GP8), and MHC-I antigen presentation (GP10) programs.

M2 regulators had generally similar impacts to those of M1, repressing MHC-I antigen presentation (GP10) and the TNF/LPS response (GP8) (albeit more weakly) and activating protein homeostasis and phagocytosis (GP5) and DC1/mDC expressed translation (GP6). Consistently, they included multiple negative regulators of NFkB and interferon signaling (E3 complex member Bid, CopA and Copb2, Ikbkg, the SUMO E3 HDAC4 and E3 Mib1 (which prevent or signal Ikba degradation, respectively), predicted E3 Nsmaf, E3 Ppp1r11 and the substrate Map3k7), and known regulators of cell differentiation in general (Bptf, Dido1, Gemin5, E3 and TF Zbtb7a), and of DC differentiation and maturation in particular (Ogt, Vdr (regulated by Mdm2), and TFs and predicted E3s Zbtb14 and Zbtb49), as well as the cell cycle and cell growth E3s APC (Anapc13, Cdc27, and Fzr1), CCNF, E4f1, Ring1, and Brca1; E3 adaptor Ddb1; E3 complex members Mdm4 and Pa2g4; DUB, Wdr48, and Fbxo11. Module members included multiple components of one complex and pathway, such as Cul4b and Ddb1 from the same cullin E3 complex (below).

M3 regulators had largely opposite effects to M2 and M1: they activated the TNF/LPS response (GP8) and autophagy and inflammation (GP9), and repressed translation (GP6) and mDCs (GP7). Module members included many known regulators of inflammation (substrate Gnb1, the E3 and TF Pparg; E3s Traf3 and Wdr82 (which regulates TRAF3)), mTOR signaling (E3s Rictor and Traf2, E3 interacting partner Mlst8; substrates Mtor, Rheb and Rptor)^74–77^, autophagy (E3 Rnf216 (Triad3); predicted E3s Pik3r4 and Wdfy3), and ER transport (E3 Syvn1, substrates Sec13 and Sec31a). In particular, the module includes Traf2, Pparg and Cebpb. CEBPB is known to activate DC2s and repress mDCs^16^, and guides targeting Traf2 and CEBPB were consistently enriched in mDCs (**Figure S1I,J**). CEBPB and PPARG physically interact^78^ and their targeting guides were enriched in CD2.2 cells. Other regulators include PAF1, which induced Tnf expression and inflammatory signaling (**Figure 2D**), and regulators of ER transport (Sec13, Sec31a, and Syvn1), all previously identified and validated as positive regulators of TNF expression in this system^5^. Our model now further relates them to mTOR pathway and autophagy regulators and to repressing mDC-like states.

M4 regulators activated ER stress (GP2), pyruvate metabolism (GP3), protein homeostasis and phagocytosis (GP5), TNF/LPS response (GP8), and DC2 MHC-II antigen presentation (GP11) and repressed oxidative stress (GP1) and translation (GP6). They included DNA repair and splicing regulators, such as the E3s Plrg1 and Prpf1 and the predicted E3 Cdc40 (Prp17), suggesting a link between DNA repair and splicing and metabolism/translational control.

M5 regulators activated autophagy and inflammation (GP9), MHC-I antigen presentation (GP10), and DC2 MHC-II antigen presentation (GP11), and repressed pyruvate metabolism (GP3), motility and cell maintenance (GP4), translation (GP6), and LPS/TNF response (GP8), through the action of regulators of antigen presentation or inflammation (SUMO E3 Pias1 and its substrate Prdm1 (negative regulators of MHC-II) and the substrates Syk, Nfkb1, and Tnf), endocytosis/trafficking (predicted E3s Wdfy2, Wdr81 and Wdr91, E3 March6), and negative regulators of CEBPB, the key DC2 TF (Rfwd2 (Cop1) and Det1 of the Cul4-Rfwd2-Det1 complex)^54, 79–81^.

Finally, M6 regulators activated mDCs (GP7) and TNF/LPS response (GP8) and repressed phagocytosis and granulation (GP1) and pyruvate metabolism (GP3), through the action of multiple cytokines and positive regulators of NF-kb (LUBAC E3s Rbck1 and Rnf31 (whose substrate Ikbkg is in “opposing module” M2), the substrate Rela; E3s Spop, Rc3h1, Traf6 and Cbl; Tlr4; and E3 complex member Bcl6), positive regulators of cytoskeleton organization and migration (RING-like Phf8^82–84^, Ub and SUMO substrate Ptpn1^85^), and E3 Keap1, a negative regulator of redox and stress that targets Nrf2^86–89^. M6 members also included multiple cullin and RING-like ligases (Cul1, Cul3, Cul5, Keap1, Rnf31, Rbck1, Brap, Arih2, Traf6), and help assign putative roles to other E3s, such as Brap, a GWAS gene for psoriasis and carotid atherosclerosis, which activated inflammatory responses in human aortic smooth muscle cells^90, 91^ but did not have a previously known cellular role in immune cells.

The co-functional modules also impacted global shifts in cell state/type distributions, consistently with the programs they regulate. For example, cells perturbed for M5 regulators were associated with a shift from naive DCs to macrophage-like cells, consistent with the module’s role as an activator of the DC2 / MHC-II presentation (GP11) program, while perturbation to M1 and M2 regulators had the opposite effect (**Figure 3B,E** and **S4C,E,F**, FDR < 0.1, Fisher’s exact test). The co-functional modules also impacted distributions within one cell state/type. For example, different modules had distinct effects on the cells within DC2.1, DC2.2 and DC2.3 subsets, (**Figure 3C,D** direction 2), orthogonally to the maturation gradient (**Figure 3C** direction 1, **Figure S4D**).

### Co-functional modules are enriched for physically interacting E3s

We next asked if our genetic regulatory network can be aligned and consistent with molecular mechanisms, such as co-complex membership, physical interactions and impact of E3s on TFs that regulate gene expression directly. First, to relate the co-functionality of E3s and complex members to shared molecular mechanisms, we searched for known protein interactions between each pair of regulators (from the STRING database^92^, **STAR Methods**) and compared those to their module membership (**Figure S4G**).

There was a significant enrichment of physical interactions between module members for four of the six co-functional modules (M1, 2, 4, and 6), as well as between one pair of modules (M3 and M5) (P<0.05, degree preserving permutation test, **STAR Methods**), suggesting that co-functional effects are congruent with joint underlying molecular mechanisms. As expected, physical interactions between members of the same (different) module were generally associated with positive (negative) correlation in functional effects (**Figure 3F** and **S4G**). Such interactions include Gbr2, Ptpn11, and Rack1 (M1), Pparg, Crebbp, Ep300, Ankfy1, and Cul2 (M3); or Cul1, Skp1a, Rnf7, Fbxw1, Rbx1, Cul3, Nedd8, Arih1, and Keap1 (M6). Within components of the NFKB signaling pathway (**Figure S4H**), multiple known TNF activators (Rela, Rbck1, Rnf31; all from M6) both physically interact with TNF and have positively correlated effects, whereas several TLR/NFKB signaling inhibitors (Nfkb1^93^, Cyld^94^, and Tab1^95^; all in M5) have physical interactions but negatively correlated effects, showing consistency between the genetic model and molecular mechanisms.

The analysis highlighted basic “rules” of co-regulation between E3 cullin, adaptor, and substrate recognition adaptor proteins, where multiple components of each Cullin complex grouped together in the same module (except the common interactor Rbx1), while the Cullin substrate recognition adaptor proteins were in other modules, consistent with their specializing or directing Cullin E3 complexes to different substrates and pathways (**Figure S4I**). For example, the SCF core complex members Cul1, Skp1 and Rbx1 are members of M6, as is Cul3, but Cul1-Skp1- Rbx1 substrate-specific adaptors or Cul3-Rbx1 adaptors are partitioned to multiple other modules (*e.g.*, Cul1-associated adaptors Fbxl14 and Fbxl5 in M2; Fbxl13 and Fbx03 in M3; Fbxw7 in M5; and Fbxo33 and Fbxw11 in M6; Cul3-associated adaptors Klhl3, Klhl24, Klhl6 in M2, M3, and M5, respectively, and Cul5-associated adaptor Socs3 in M5; **Figure S4I**). Similarly, Cul4b and its adaptor Ddb1 are part of M2, but Dda1, which recruits and organizes substrate receptors with both Cul3 and Cul4 complexes, is in M6. Furthermore, each of the interacting pairs of Rbx1 and Arih1 and Cul5 and Arih2 are in M6, consistent with their physical interaction and known functional roles in forming highly specific neddylated CRLs^57^, but their expected adaptors were not (e.g., most F-box proteins for Cul1 and Arih1; except Fbxw7 and 11). This highlights the versatility and modularity of the ubiquitin system, whereby different adaptors/receptors target different substrates regulating different aspects of DC lifecycle, and how a systematic Perturb-Seq screen and computational analysis can decipher this organization.

### E3 perturbation effects on gene programs explained by modulation of TF activities

Next, we examined how the E3 ligases may propagate to the transcriptional level, by combining our genetic model with one associating TFs to their physical targets, to infer TFs whose activity (as inferred from the expression of its known targets^96, 97^) is impacted by each perturbation (**STAR Methods**).

Overall, the 329 knockout (KO) perturbations were significantly associated with inferred changes in activity of 123 TFs (**Figure S4J**), with 32 TFs with the most prominent effects from 41 E3 ligase and complex members (**Figure 3G**). Most notably, Cul3 and Keap1 perturbations decreased the activity of many TFs, including Nfkb1, Nfkb2, Rela, Relb, Jund, Irf1, Cebpb, Nfya, Klf4, Foxo3, and Foxo4 and increased the (inferred) activity of just four of the 32 highly regulated TFs: Atf6, Wt1, Srf, and Pax6 (**Figure 3G** box 1). Importantly, we show Cul3 and Keap1 perturbations increased (inferred) activity of the well known CUL3-KEAP1 substrate and master regulator of oxidative stress, Nrf2 (**Figure S4J**). The set of TFs whose activity was impacted by Cul3 and Keap1 perturbations was further partitioned to two subsets based on their opposing regulation by different adaptors and receptors. For example, Rack1, Dcaf13 and Vprbp perturbations increased the (inferred) activity of Foxo3/4, Klf4, Twist1, and Smad1 (**Figure 3G**, box 2), while perturbations of M6 E3s, including Spop, Traf6, Fbxw7, Fbxw11, and Cul1 decreased the (inferred) activity of Irf1, Nfkb2, Nfkb1, Rela, Relb, Rel, and Jun (**Figure 3G**, box 3).

This analysis detects established links between E3s and regulated TFs, supporting its validity. For example, Hif1a activity increased following KO of any member of the E3 ligase complex VHL-TCEB1-TCEB2, which binds and ubiquitylates Hif1a for degradation^98^ (**Figure S4J**). Cebpb and Jun’s inferred activity increased in cells with KO of the E3 ligase Rfwd2 (**Figure 3G**), a member of the CUL4-DDB1-DET1-RFWD2 complex that targets Cebp family TFs and Jun for ubiquitination and degradation^80^. Indeed, Rfwd2 represses programs that are activated by Cebpb (GP8) or Jun (GP1 and 8) and are enriched for their bound targets (**Figure S4K**). Because Rfwd2 knockout similarly increases the (inferred) activity of Foxl2 and JunD, these could be additional Rfwd2 substrates.

By relating joint TF targets, E3 targets and E3 regulated programs, we explain E3s impact on different programs through different mediating TFs. For example, by this analysis, Rela mediates Cul3 and Keap1 effects on GP7 and GP8, but not on GP1 (**Figure 3H**), and E3 Fbxw11’s impacts on GP7 (but not GP4) (**Figure 3I**). Notably, Cul3, Keap1, Fxbw11 and Rela itself are all members of M6, and the expression of Rela’s targets in GP7 is decreased when any of these regulators is perturbed (KO) in our screen (**Table S8**; **Figure 2C,D**). However, because KO of Cul1, Keap1 and Fbxw11 leads to (inferred) reduction in activity of Rela, it is unlikely that Cul3, Keap1 or Fbxw11’s effects is through targeting of Rela for degradation (see **Discussion**).

### Perturbed E3 ligases impact multiple statistically-independent pathways

The effect of perturbing one gene on another gene’s expression can be due to various pathways with direct or indirect dependencies and indeed pairs of programs in our regulatory model are dependent, as reflected by their pair-wise positive and negative correlations (**Figure 2C**). To decompose the observed regulatory effects into a set of statistically-independent factors, we performed Independent Components Analysis (ICA)^99^ on the regulatory matrix, recovering 15 independent latent factors whose weighted sums optimally explained the perturbation effects (**Figure S5A-E, STAR Methods**), and annotated them by enrichment in functional gene sets and known marker genes (**Table S10**). On average, the factors explained 35% of the variance of observed effects of the 329 impactful regulators (27 explained well (>70%); 72 poorly (<20%), **Figure 4A**, bottom, **STAR METHODS**). The latent factors have little to no correlation (maximal Spearman ρ = 0.14) and little overlap in highly loading genes (**Figure S5B,G,** maximum Jaccard similarity index = 0.09), but perturbed genes can affect multiple independent factors (**Figure S5A,H**, maximum Jaccard similarity index = 0.4).

**Figure 4.**
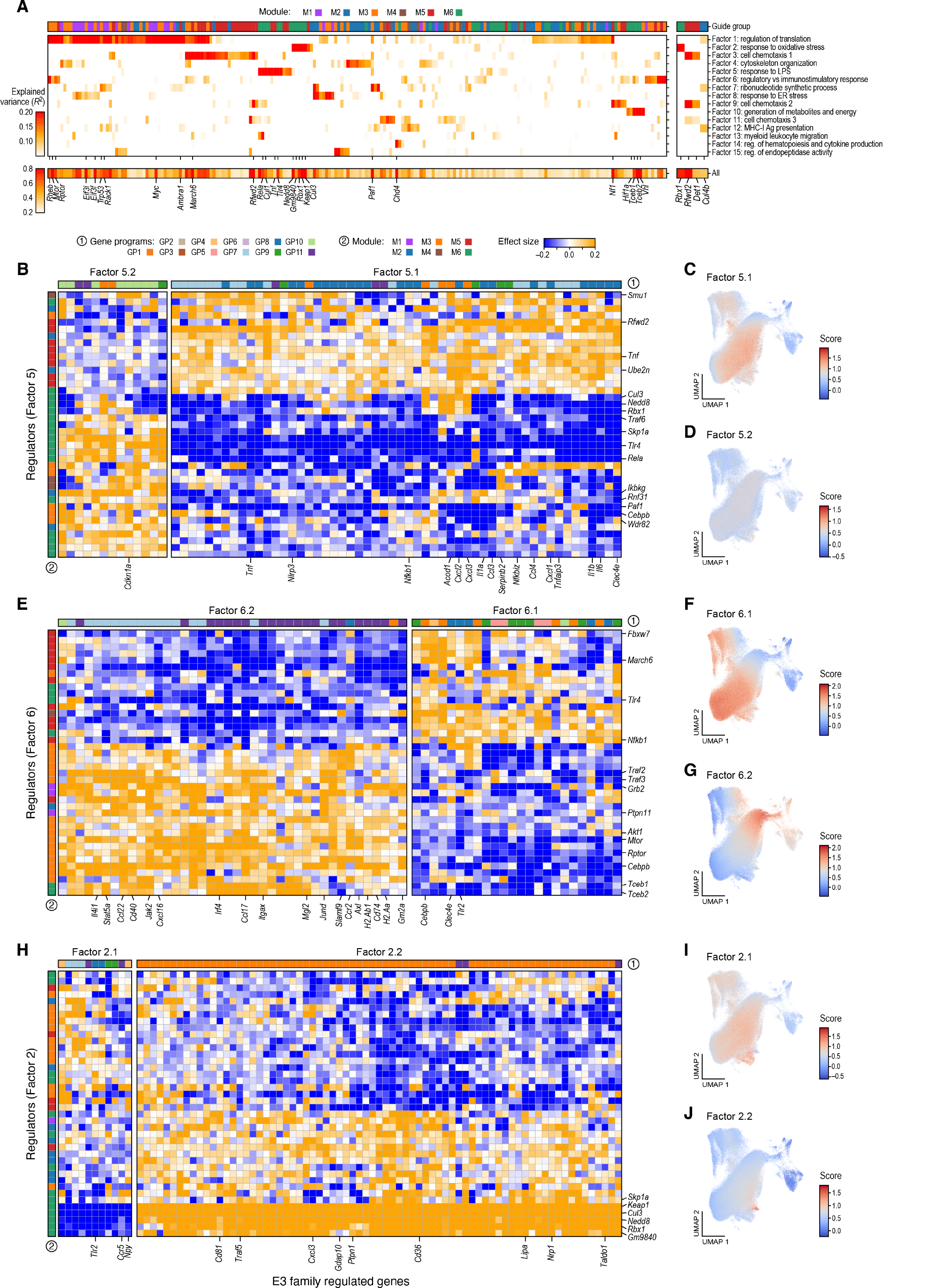
Independent programs affected by distinct combinations of E3-ligase family members. **A.** E3 regulators association with independent factors from ICA. Left: Explained variance of effects on 1,041 genes of each of 203 perturbed genes (left matrix columns, sum of explained variance > 25%) or components of the CUL4-RBX1-DET1-RFWD2 complex (right matrix columns) by each of 15 latent factors (rows, main panel) and across all 15 factors (bottom). **B-J.** Member genes and regulators of example ICA factors. **B,E,H.** Effect sizes (yellow: positive; blue: negative; color bar) on significantly affected genes (columns; outlier loadings, separated by direction of effect) upon perturbation of each regulator gene associated with the factor (rows; outliers based on weights in the mixing matrix) (**STAR Methods**). Left bar: Co-functional modules; top bar: gene programs from the regulatory model. **C,D,F,G,I,J.** UMAP embedding of cell profiles (as in Figure 1C) colored by expression scores for each sub-factor (label on top). See also related **Figure S5**.

Each factor simultaneously captures both induced and repressed genes along with diametrically opposed regulators. For example, the LPS response factor (#5, **Figure 4B-D**) includes activation of LPS and TNF response genes and repression of genes more highly expressed in DC2s *vs*. mDCs and DC1s (**Figure 4C,D**). It is positively regulated by well-established activators of TNF and the LPS response (*e.g.*, Rnf31, Traf6, Paf1, Ikbkg, Rela, Cebpb) ^5^, and repressed by known negative regulators (*e.g.*. Rfwd2^54^), as well as by previously-uncharacterized regulators, both positively (E3 adaptor Skp1a) and negatively (E2 Ube2n; E3 substrate adaptor Smu1). The DC immune control factor (#6, **Figure 4E-G**) consists of inflammation and cytokine response genes in two opposing patterns, capturing the maturation gradient from immunostimulatory monocyte derived macrophages and DC2s to immunomodulatory mDCs genes, and its corresponding, diametrically-opposed regulation by E3 Traf2 and E3 adapter Ptpn11 *vs.* March6 and Fbxw7, respectively.

Because one regulator can be associated with multiple ICA factors, this decomposition groups together different subsets of multi-subunit E3 complex members. For example, all four components of the Cul3-Skp1a-Rbx1-Nedd8 complex are associated as negative regulators of the response to oxidative stress factor (#2) (**Figure 4H-J**). Conversely, different combinations of subunits of one complex are associated with different ICA factors, predicting how interactions of one core complex with different substrate recognition adaptor proteins can drive a variety of gene regulation programs (as also described above). For example, in the Rfwd2/Cul4a/Ddb1/Rbx1/Det1 complex, Rfwd2 (Cop1), the E3 substrate recognition adaptor protein that forms an active E3 complex, has outlier loadings indicating regulation of ICA Factors 3, 5, 9, 10, and 11 (**Table S10**). Other members of this complex regulate other factors, not impacted by Rfwd2 perturbation (*e.g.*, Ddb1 regulates Factor 2, **Table S10**). Interestingly, while Rfwd2 and Det1 knockout are both similarly strongly associated with the same factors (#3, 9, and 11), other complex members knockouts (Rbx1, Cul4b) have only weak associations in those same factors (**Figure 4A**, right panel). Thus, Rfwd2 or Det1 may interact with other E3 or cullin complex members, just as the Cul4 complex interacts with other substrate recognition adaptor proteins.

### Intra-module genetic interactions and inter-module additivity in combinatorially- perturbed cells

In the regulatory network, many of the E3s and other regulators impact the same genes and processes when perturbed individually, but, given the possible non-additivity of biological interactions, determining their effect if perturbed jointly (combinatorial perturbation) requires an additional experiment. In our large screen, 177,871 cells had more than one guide assigned (with 10,244 cells with guides targeting two or more of the 329 singly-impactful regulators; **Figure S6A**), opening the way to test for such genetic interactions. However, because of random sampling from an enormous number of possible combinations, few to no cells were profiled for any specific combination, such that we could not determine the joint effect of any individual combination with confidence. Instead, we reasoned that we can leverage the co-functional modules to group all cells perturbed by a pair of guides from a given pair of modules (including the same module) to gain statistical power to decipher genetic interactions between or within modules rather than between their individual constituent genes. We thus analyzed double perturbations in a setting where cells are assigned to perturbed modules instead of perturbed genes. (The few detected triply perturbed cells were removed prior to analysis, as were module 4 perturbations given the very small number of cells.)

The proportion of genes whose expression has a significant interaction term due to a combinatorial perturbation was much greater in intra- *vs*. inter-module combinations (**Figure 5A**). Thus, when two genes within the same module (intra-module combination) were perturbed in the same cell, the combined effects on the expression of genes were often different than the sum, with a super-linear relationship in M1, M2, and M6 (**Figure 5A,B, STAR METHODS**). Conversely, when two genes from different modules were perturbed in the same cell (inter-module combinations), the impact on most genes was additive (**Figure 5A**).

**Figure 5.**
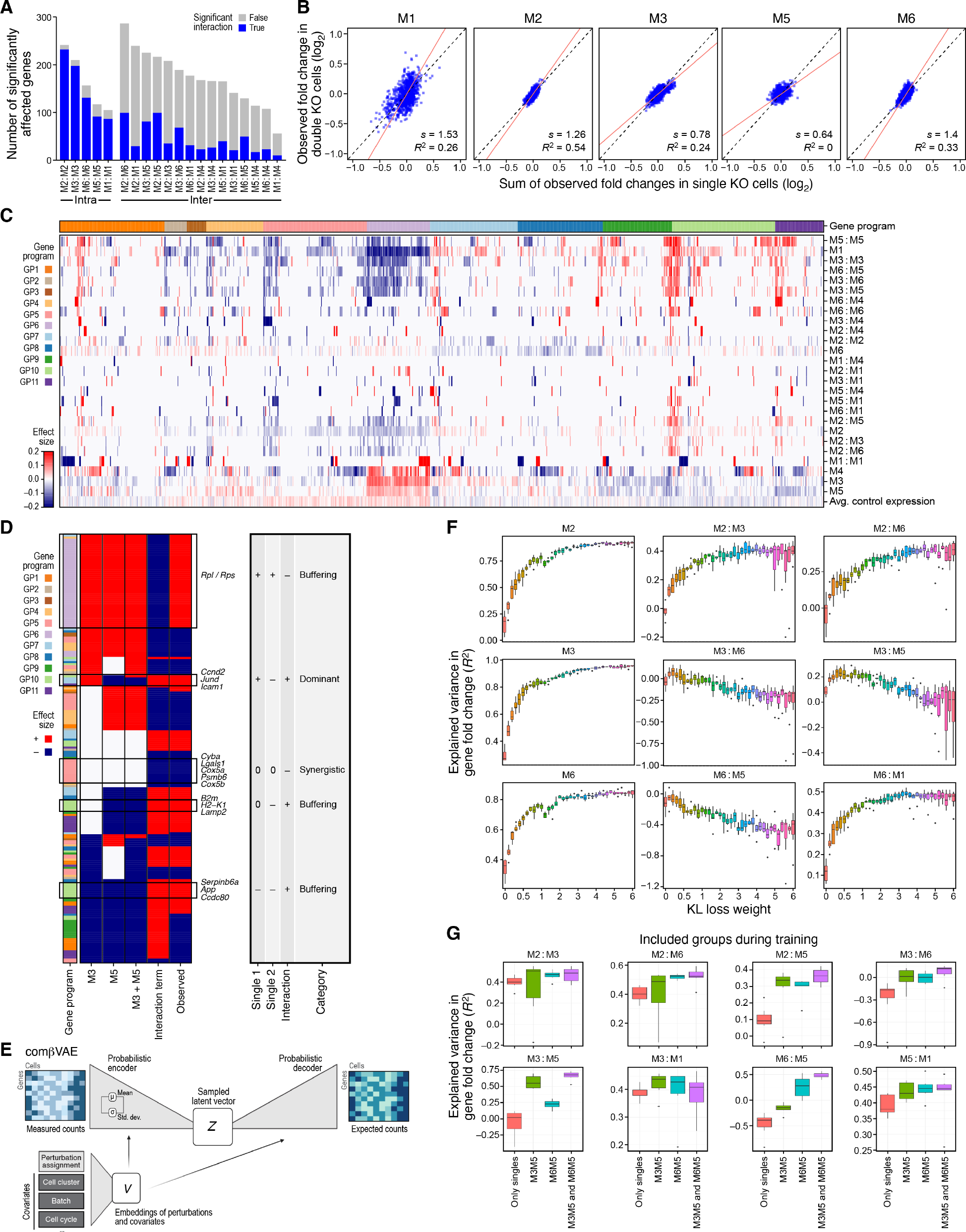
Intra-module genetic interactions are more prevalent than inter-module interactions and modular structure can be leveraged to predict combinatorial perturbations **A.** Inter-module interactions are more prominent. Number of genes (y-axis) significantly affected by intra- (left bar) or inter- (right bars) module combinatorial perturbation (x-axis), additively (grey) or non-additively (blue). **B.** Intra-module interactions. Observed (y axis) fold-changes in expression (*vs*. control) in cells with perturbations in two genes from the same module and the expected fold change (x axis) from an additive model based on the two individual perturbations for each of 1,041 genes (dots). Slope of the first PC (red line) and variance in observed double knockouts explained by the single knockouts (R^2^) are labeled. Module M4 is not shown to insufficient number of double-knockout cells. **C.** Inter- and intra-module interactions vary across programs. Significant effect sizes (red/blue color bar; FDR<0.1) of perturbations at the level of individual modules (M_i_), inter-module pairs (M_i_:M_j_; i≠j), and intra-module pairs (M_i_:M_i_) (rows) on each of 1,041 genes (columns, labeled by gene program). Bottom row: Row centered mean expression in control cells. **D.** Substantial interactions between M3 and M5 regulators. Binarized significant effects (FDR<0.1, red/blue: positive/negative) on gene expression (rows, only genes with significant interaction terms) by single perturbations in regulators from M3 or M5, their additive effect (M3+M5), their interaction term (M3:M5), and observed combined effect (columns). **E-G**. comβVAE predicts combinatorial perturbations. **E.** Method overview. **F.** Distribution of explained variance (R^2^, y axis) in fold changes of the 1,041 genes from 7 runs with the same hyperparameters at different KL loss weights (x-axis) for individual modules (M_i_) and inter-module combinations (M_i_:M_j_; i≠j). **G.** Distribution of the explained variance (R^2^, y axis) in fold changes of the 1,041 genes from 7 runs with the same hyperparameters in the indicated inter-module combinations (labels on top) when the model (Beta=6.0) is trained only with data from single KOs from all modules (M) or single KOs from all modules and double KOs from one or two pairs of modules (M_i_:M_j_; i≠j) (x axis). Boxes display the first (Q_1_), second (Q_2,_ median) and third (Q_3_) quartiles while the bottom and top whiskers show the intervals [Q_1_ -1.5 IQR, Q_1_] and [Q_3,_ Q_3_ +1.5 IQR], respectively. See also related Figures S6 and S7.

At the level of expression of the individual affected genes, we found a range of patterns, with almost half (490 of 1,041 tested genes) with at least one significant interaction term (either positive (synergistic) or negative (antagonistic)) as a result of at least one of the 15 inter-module KO pair groups, and 650 with interaction terms in the five intra-module perturbation pairs (FDR < 0.1, **Figure 5C,D** and **S6B,D,E**, **STAR METHODS**). In particular, a joint perturbation of an M3 and M5 regulator yielded non-additive effects in many genes (**Figure 5D**), enriched for biosynthetic, translation, cytokine production, and inflammatory response genes. These included buffering for translation (GP6) and MHC-I presentation (GP10) program genes; synergy for protein homeostasis and phagocytosis (GP5) program genes, and dominance for mDC (GP7) program genes (**Figure 5D**). Notably, the impacts of single perturbations in M3 and M5 regulators are often correlated (**Figure 2C** right and **5D**). Thus, similarly to the joint perturbation of two regulators from the same module, non-additive effects may be more prevalent for regulators from different modules but with similar effects on gene programs when individually perturbed.

### comβVAE predict combinatorial perturbations within and across modules

We next asked how well the effects of the double knockouts can be predicted from profiles of single knockout cells. As a baseline, we first used a simple linear model (**STAR METHODS**) to assess the overall effects of each of the six co-functional modules on the 1,041 response genes, and predict the log_2_ fold changes of these genes in 20 pairwise module combinations by adding the individual KO group effects. As expected from our analysis above, additive effects explained most of the intra-module interactions quite poorly (**Figure 5B**), while a substantial fraction of the variance in fold changes was explained for some of the inter-module pair combinations (**Figure S6F-H**). Inter-module groups where the additive model performed worse involved pairs of perturbations where more target genes show significant interactions, such as M2-M5 and M3-M5 (**Figure 5A** and **Figure S6C,F-H**), or those with fewer double KO cells, like M1-M4 (**Figure S6A,F-H**), which may have affected the estimation of the ground truth values. For genes with significant interaction terms, the additive model lost its predictive power in most of the module pairs (**Figure S6F-H**), including a change in effect direction for many genes between prediction and observation (**Table S11**).

We hypothesized that we could gain better prediction performance by learning the interaction effects based on the latent structure of the gene expression profiles. To test this hypothesis, we developed com VAE, a conditional variational autoencoder (CVAE)^100, 101^, where the latent variables of the observed data are distributed conditioned on input data labels (**Figure 5E**, **STAR METHODS**). We trained our model with 89,463 single KO cells (of 329 impactful KOs, 80,189 cells for training, 9,274 cells for validation) and a random sample of 70% of control cells, conditioning on 7 groups (controls + 6 regulator modules). With the remaining 30% of control cells, we used the trained model to generate profiles based on the counterfactual questions “*What would be the profile of this control cell if it had a single knockout from module x*?” and “*What would be the profile of this control cell if it had a double knockout, one from module x and another from module y*?” Note that this model only addresses inter-module interactions, as the conditioning is done per KO module (**STAR Methods**). Finally, we calculated the expression fold changes between these generated cells and the population of control cells.

While the explained variance in the expression fold changes for generated KO profiles of single genes from the groups observed during training (single knockout modules) was quite high (0.77<*r*^2^<0.95, mean 0.85), estimates for double knockouts varied based on the module pair (**Figure S7A,B**). The profiles of cells with pairs of KOs from M3*M5, the module pair with the highest number of significant inter-module interactions (**Figure S6C**), were estimated far better by com VAE (*r*^2^=0.23) than by the additive model (*r*^2^=0.02) (**Figure S6F, S7A,B**). Moreover, com VAE had smaller mean absolute errors than the additive model when predicting the fold changes of genes with significant interaction terms (FDR< 0.1), especially in module pairs with the highest number of genes with significant interactions (M3*M5, M3*M6 and M6*M5; **Figure S6G, S7C**). Higher values of Beta, the hyperparameter that changes the weight of the Kullback-Leibler (KL) loss term (which acts as a regularizer on the latent space distribution of the gene expression embeddings as well as the KO embeddings, **STAR Methods**), increased the explained variance in single KOs modules and in double KO module pairs with fewer genes with interaction terms, but reduced it for pairs with more genes with interaction terms (**Figure 5F** and **S6C, S7D**). Thus, greater latent space disentanglement (*i.e.*, larger regularization parameter Beta) leads to better conditioning on the individual single KO groups, and better learning of additive effects at the cost of non-additive ones.

To evaluate how the predictions change when we include some double KO cells during training, we trained our model with the training set of the singly perturbed cells and the double KO cells of either M3M5, M5M6, or both. Interestingly, including double KO cells from one pair of modules increased the prediction performance of *other* unseen double KO groups (**Figure 5G** and **S7E**). Furthermore, when double KO cells were included in the training, higher beta values increased the prediction performance of inter-module groups with more interaction terms (**Figure S7F,G, Table S11**). Thus, including some combinatorial perturbations during training along with better latent space disentanglement helps the model to learn both the generative factors and connections between them, leading to better prediction of unseen combinations.

### *In vitro* perturbation-defined regulators and programs are associated with genetic risk in inflammatory diseases

To determine if our regulatory network contributes to or is active in human disease, we asked whether impactful regulators in the modules or the programs they regulate are also likely to be causal in human disease, based on either heritability signals, regulation *in vivo* during disease progression, or both. We thus tested the modules and programs for enrichment in common-variant driven disease associations using sc-linker^102^ and MAGMA^103^ (**STAR Methods**) with GWAS summary statistics from nine immune-related diseases (average N = 79.5K, **Figure 6A,B** and **Table S12,S13**). We also compared each gene program to cell type specific disease progression programs induced in disease *vs*. healthy tissue by scRNA-Seq^102^ (**Figure 6C,D**, **STAR Methods**). We found support for both relationships.

**Figure 6.**
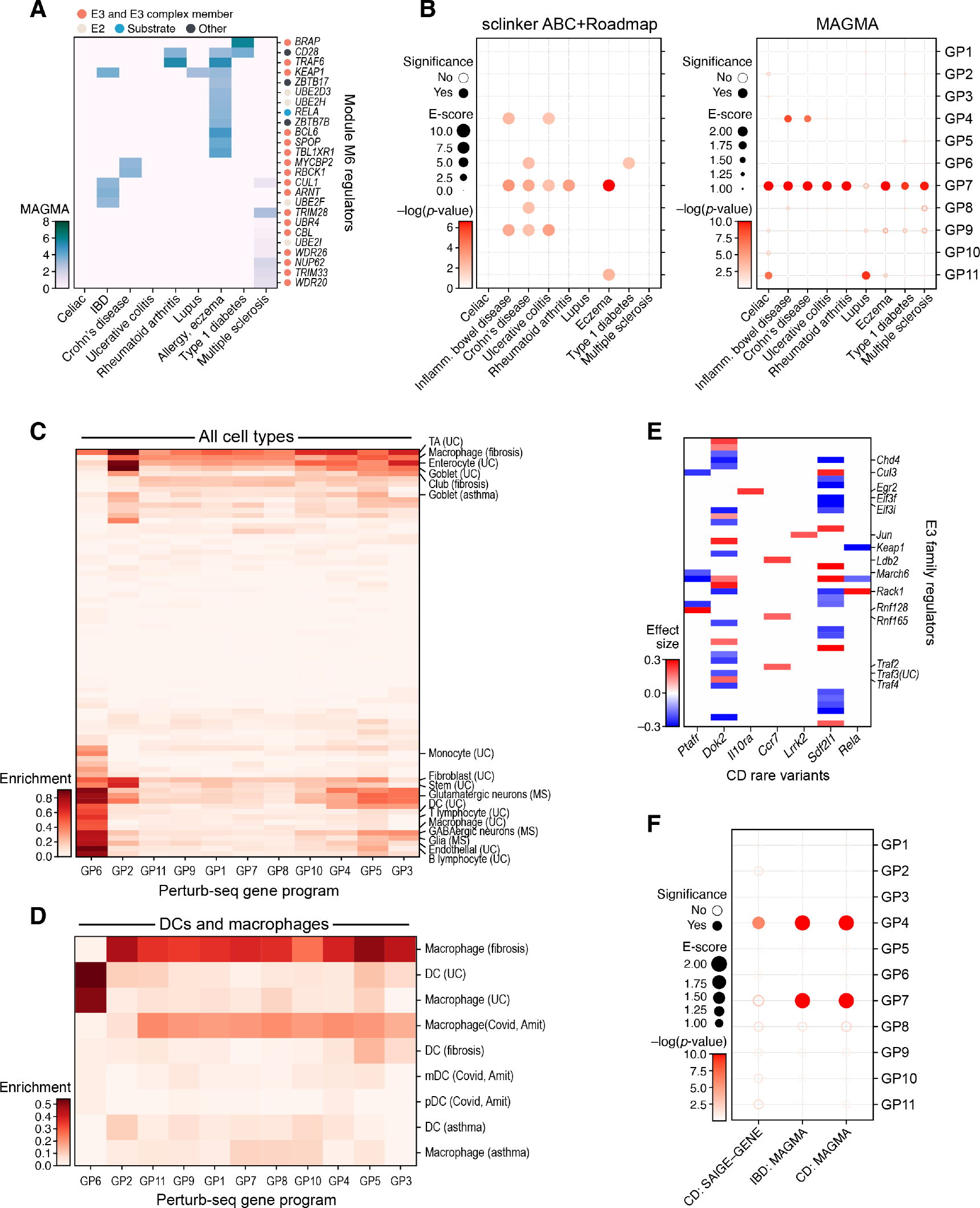
Co-functional modules and gene programs are enriched for risk heritability and cell programs in human immunological disease. **A.** Module M6 associated with genetic disease risk for human immunological disease. Significant MAGMA Z-scores (color bar; per trait; Bonferroni α < 0.1) for immunological disease traits (columns) of module M6 E3 family genes (rows) with at least one significant score. **B.** Gene programs associated with genetic disease risk for human immunological disease. Significance (-log_10_(p-value), dot color) and effect size (dot size) of heritability enrichment in each gene program (rows) for different immune traits (columns) by sc-linker analysis with SNP annotations combined with intersection of Roadmap and ABC gene-enhancer linking strategy (left) or by MAGMA (right). **C,D.** Genes programs expressed during immunological disease progression in immune and non-immune cells. Enrichment (color bar) of gene programs (columns) for cell type specific disease progression programs in humans (rows), across diverse cell types and diseases (C) or only in DCs and macrophages in UC, fibrosis, asthma, and COVID-19 (D). **E.** Perturb-Seq regulatory model highlights E3 regulators of rare IBD disease genes. Regulatory coefficient (color bar, from the model of Figure 2C) of the impact of perturbing regulators (rows) on the expression of genes (columns) with rare variants associated with IBD that also have at least one significantly regulating E3 family member. **F.** Gene program enriched for rare variant in immunological disease. Significance (-log_10_(p-value), dot color) and effect size (dot size) of heritability enrichment of gene programs (rows) for CD or IBD based on rare (CD:SAIGE-GENE) or common (IBD:MAGMA and CD: MAGMA) variants.

Among the co-functional modules, common and low frequency variants in genes in module M1 were enriched for heritability across all traits tested (1.62-fold on average, P = 1.52x10^-5^), and especially immune-related traits (1.94 fold; P = 2X10^-4^), compared to genes constituting all the modules (**Table S12**). In particular, M1 member gene and predicted E3 *WDR36* has a significant MAGMA score in allergy/eczema and blood traits and E3 adapter *PTPN11* in type 1 diabetes (T1D) and blood traits (Z-scored per-trait MAGMA scores, Bonferroni correction α = 0.1). Notably, perturbing module M1 activates the mDC (GP7), TNF/LPS response (GP8), and MHC-I Ag presentation (GP10) programs and represses ER stress (GP2), protein homeostasis and phagocytosis (GP5) and translation (GP6) (**Figure 2C,D** and **Table S13**), consistent with the association of variants in this regulating module with inflammatory disease. Additionally, many E3s and E3 complex members in M6 are associated with risk of immune-related traits (**Figure 6A**), including Traf6 (eczema and RA), Rbck1 (CD), Bcl6 (eczema), Keap1 (eczema, IBD, and lupus), Brap (T1D), and Cul1(IBD) (Z-scored per-trait MAGMA scores, Bonferroni MTC a = 0.1; **Figure 6A**). Several co-regulated gene programs also showed heritability enrichment in immune diseases. These included the mDC program (GP7) (**Figure 6B**, highest score by both sc-linker and MAGMA), especially *REL* (rank 1 average MAGMA scores across immune diseases), *JAK2* (rank 3) and *STAT5A* (rank 8); and the motility and cel1 maintenance (GP4) program in IBD and related traits (**Figure 6B**), with top driving genes *CCL2* (rank 2) and *C*CL7 (rank 9)^104, 105^.

Concomitantly, several of the perturbation-affected programs were enriched (relative to all genes expressed in this cell type) in disease progression programs from multiple cell types and diseases, especially those of three key cellular processes that are also implicated in modulating immune responses – mitochondrial metabolism^26, 106^, ER stress^107^, and translation/antigen presentation. This is consistent with a model where processes affected by heritable variation in regulators lead to dysregulation of their target programs. For example, translation program genes (GP6) were enriched in disease progression programs of immune and non-immune cells in inflammatory diseases, including ulcerative colitis (UC) and multiple sclerosis (MS) (**Figure 6C,D**); ER stress response genes (GP2) were enriched in disease progression programs in epithelial cells in UC, lung fibrosis and asthma (**Figure 6C**); and disease progression programs in macrophages and DCs in UC are enriched for the translation program (GP6) (**Figure 6D**). These programs are all regulated by module M1, which is itself enriched for heritability of disease risk, as noted above. Thus, our analysis suggests that at least two modules (M1 and M6) may contribute to disease risk through their respective activation or repression of disease induced programs including ER stress (GP2), motility and cell maintenance (GP4), and translation (GP6), including by E3 risk genes Wdr36 (M1) and Keap1, Cul1, and Rbck1 (M6). In addition, the GP4 program is enriched for genes with rare-variant driven association in Crohn’s disease^108^, including *COX4I1*, *POLD4* and *NPY* (ranked 1, 2 and 4; **Figure 6F**). Interaction between *NPY* neurons and immune cells in the enteric nervous system has previously been implicated in IBD pathogenesis, and *NPY* expression changes occur in animal models of IBD^109, 110^.

Finally, we utilized our model as a “look up” resource for proposing regulators of rare risk variants for Crohn’s disease^108^. Our model suggests that Il10ra expression is repressed by Egr2 and that expression of Ccr7, a chemokine receptor regulating many aspects of DC function and guiding DCs to lymph nodes^111^, is repressed by Ldb2, Traf2, and Rnf165 (**Figure 6E**). While Traf2 deletion impacts inflammation in both DCs and keratinocytes^112^, Traf2-based regulation of Ccr7 has not been previously reported. Increased Ccr7 expression could be an important mechanism in increased T cell infiltration and inflammation controlled by Traf2.

Thus, multiple components of our model, including regulatory modules and their impacted programs are congruent with both risk genes for human immune and inflammatory disease and the dysregulated expression programs observed in patients. This highlights the relevance of our *in vitro* screen to human disease.

## DISCUSSION

In this study, we characterized a large gene family, the E3 ligases, and their interacting partners in the cellular response of primary immune cells to an inflammatory signal. We showed the power of systematic Perturb-Seq to relate different members of one gene family as regulators in distinct co-functional modules, and their impact on individual genes, co-regulated gene programs, and cell state distributions, across a mixed population of related cell types. The modular organization of the regulatory network further allowed us to study and predict the impact of genetic interactions and relate *in vitro* perturbations in a model system to the mechanisms underlying disease risk in humans.

### E3s regulate key phases of the DC life cycle

The independent factors and programs of target genes regulated by perturbations in E3s and associated proteins span multiple stages in DC life cycle (**Figure 7**), showing the capacity of our screen to capture many phenotypes, and the breadth of roles E3s play in the immune response and the DC life cycle, many of which are novel roles. These include, in order of the lifecycle, regulation of: (**1**) differentiation towards DC2s (*e.g.*, Cul3- Keap1), DC1 state (*e.g.*, Arih2), and mDCs (*e.g.*, Traf2); (**2**) sensing, including the response to LPS (factor #5), ER stress (#8), and oxidative stress (#2), ribonucleotide synthesis (#7), and metabolism and energy (#10), regulated by *e.g.*, the Keap1-Cul3-Rbx1 complex, the Tceb1-Tceb2-Rbx1-Vhl complex, and Traf6; (**3**) DC migration, including chemotaxis (#3,9,11), cytoskeleton organization (#4), and myeloid migration (#13), regulated by *e.g.*, Pparg, Rfwd2, Brap, and the Keap1-Cul3-Rbx1 complex; (**4**) antigen presentation and associated processes (antigen presentation (#12), endopeptidases (#15), and translation (#1)), regulated by *e.g.*, Ambra1, March6, Traf2, and Traf3; and (**5**) production of chemokines that promote either a regulatory or immunostimulatory response (#6.1, #6.2, #14), regulated by *e.g.*, Traf2, Traf3, March6, Pias1, Plgr1, Ptpn11, and Prpf19.

**Figure 7.**
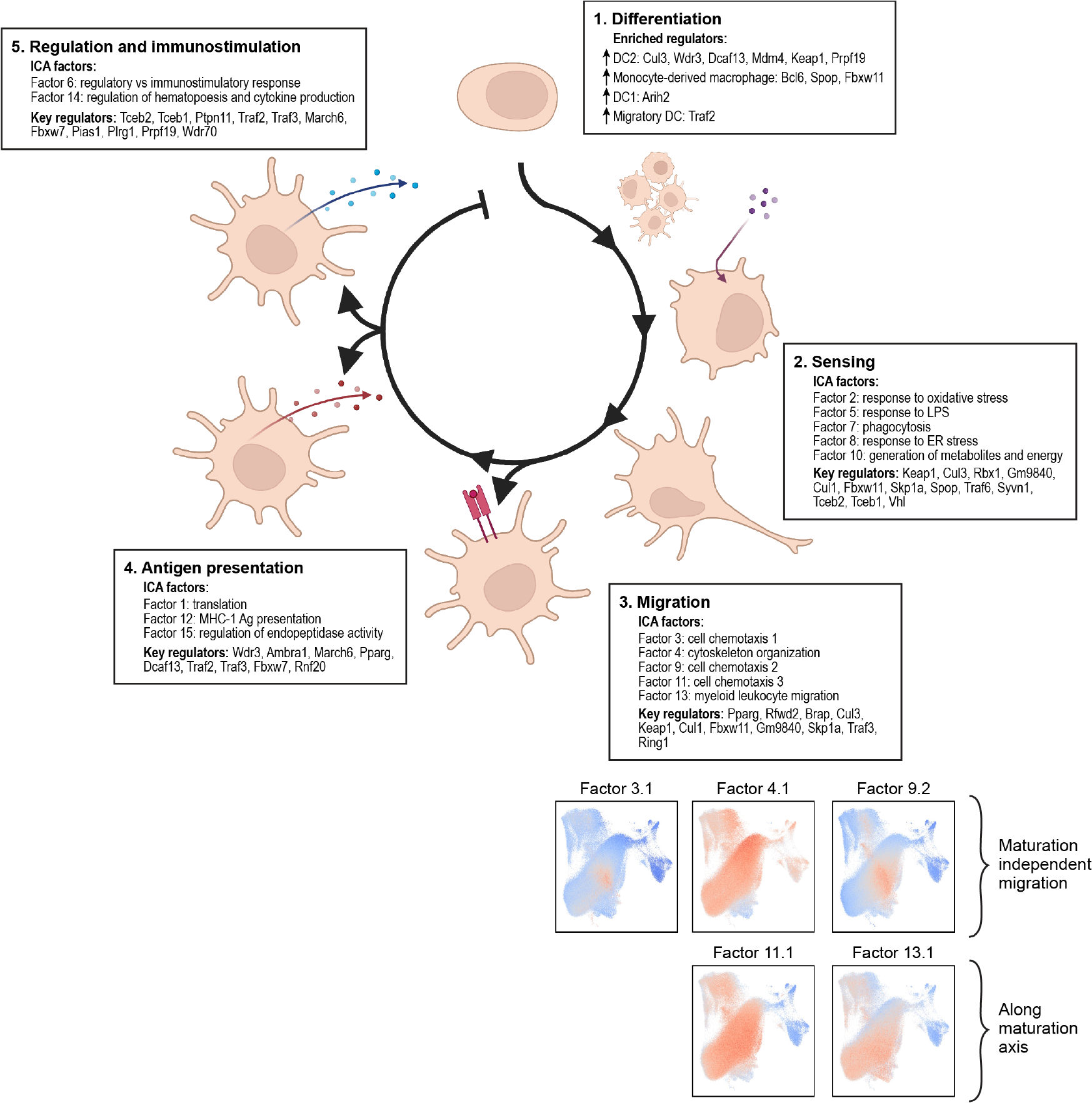
The DC life cycle regulated by E3 ligases ICA factors and their key regulators grouped in each DC lifecycle stages (boxed). Bottom: UMAP embedding of cell profiles (as in Figure 1C) colored by expression scores (color bar) for migration-related factors.

The detailed regulatory model highlights many novel regulatory relations and helps address open questions. For example, it has been unclear whether DC maturation and migration are inextricably linked^113, 114^. Our model shows that while the expression of some migration factors (#11; #13) follow a cell maturation gradient, other factors (#3, #4, and #9) express migration genes independent of DC maturation. While the same regulators are shared across migration and maturation in some factors (*e.g.*, Cul3-Keap1 and the CLR1 complex which co-regulate in Factor #4), they regulate them in opposite ways in others (*e.g.*, Cul3-Keap1 and the CLR1 complex in Factor # 9 or #13). This suggests that some, but not all, of the DC migration program is controlled independently of maturation. Several E3s are regulators of multiple programs along the DC lifecycle (*e.g.*, Keap1-Cul3-Rbx1, Fbxw11-Cul1-Skp1a, Fbxw7, March6), while others play specific roles in key stages (*e.g.*, Rfwd2 (Cop1) in migration; Wdr70 in immunostimulatory *vs*. immunoregulatory response).

### Congruent genetic effects and physical interactions relate complex members and characterize E3 partners and substrates

Screening both E3s and their interacting partners and substrates allowed us to relate functional (genetic) effects to physical interactions and molecular mechanisms. Co-functional modules of regulators are enriched for physical protein-protein interactions and members from the same E3 complex. While multiple components of each Cullin complex grouped together in the same module (except the common interactor Rbx1), the Cullin substrate recognition adaptor proteins were in other modules, highlighting their specializing or directing Cullin E3 complexes to different substrates and pathways. Because programs may not be independent, and regulators are strictly partitioned to separate modules, when regulators have multiple roles, this partitioning can mask their full set of relationships. For example, in the CUL4-DDB1-RBX1-DET1-RFWD2 E3 cullin complex, Cul4b and its adaptor Ddb1 are both members of M2, and Rbx1, Cul1/3/5, and Det1 are in M5. This challenge is addressed when considering independent factors associated with overlapping regulators: Rfwd2 (Cop1) co-regulated the response to LPS with Rbx1 (factor #5), the response to oxidative stress with Ddb1 (#2), and chemotaxis with Det1 (#9). Thus, the ICA factors allow us to relate different adaptors with partly overlapping effects (*e.g.*, Rbx1, Det1, and Ddb1) and the way in which they combine with different substrate recognition adaptor proteins.

Combining co-functional genetic profiles with physical interactions showed congruence between genetic relations and molecular mechanisms, and helped suggest new interactions between E3s and putative adaptors. For example, surprisingly, Rfwd2, but not other members of the CUL4- DDB1-RBX1-DET1-RFWD2 complex, is a regulator of factors 3, 10, and 11, suggesting that it may interact with other complexes to ubiquitylate targets. Other regulators of these factors that have strong physical interactions with Rfwd2 and could be such candidates include: Ptpn11 (Shp2) (factor 3 and 10 regulator) that acts as an adaptor with p38-pRfwd2 to bind and catalyze Ub-mediated degradation of FASN^115^; Wdr82 and Ep300 (factor 10 and 11); Anapc13^116^, Cul2, Cul5, and Huwe1 (factor 10); and Crebbp, Skp1a, Nedd8, Cul1, and Wdr5 (factor 11).

Because knockouts of E3s and their substrates should have opposite effects when the substrate is targeted for degradation, and similar effects when Ub modification is activating, we can predict the directionality of protein level regulation by the correlation between expression profiles of E3-substrate pairs in our screen. For example, the regulatory profile of Cebpb is negatively correlated with that of Rfwd2 in all ICA factors where both are regulators (#3,5,11), consistent with the targeting of CEBPB for degradation by the CUL4-DDB1-RBX1-DET1-RFWD2 complex. Conversely, Fbxw11 KO leads to repression of (inferred) activity of all of NFKB1 (p105/p50 precursor), NFKB2 (p100/p52 precursor), Rela (p65) and Relb. Because the targets of these transcriptional activators are repressed both by the TFs own KO and by Cul3, Keap1 and Fxbw11 KO, it is unlikely that this effect is mediated by the TF’s degradation. For Cul3 and Keap1, the effect is likely through direct CUL3-KEAP1 ubiquitin modification and subsequent degradation of IKBKB^117^. As for Fxbw11, it is reported to directly bind with NFKB1 and NFKB2^118^, and this binding is enhanced in the presence of a proteasome inhibitor^119^. While the current model suggests that full length Nfkb1 is processed constitutively and Fbxw11 (also known as BTrCP2) targets Nfkb2 for complete degradation upon stimulation such as LPS activation^118, 120^, our data may not be fully consistent with such a model. We hypothesize that Fbxw11 could be part of the system that interacts with the proteasome to processes p105 (Nfkb1) and p100 (Nfkb2) to generate active p50 and p52, respectively. These analyses may be particularly helpful for multi-subunit E3 Ligase complexes reusing core scaffolds and adaptors with different substrate recognition adaptor proteins.

### Scaled genetic perturbation and combinatorial screens

We leveraged several efficiencies to enable Perturb-seq at scale, including hashing and overloading (40,000 cells per droplet channel; 5-fold cost reduction) and shallow sequencing (15,900 reads per cell on average; 2-fold reduction). Future efficiencies could include pre-barcoding (for even higher overloading^121^), cheaper sequencing^122^, and further guide-compressed screens^123^,(Yao et al, bioRxiv 2023). For guide compression we note that we initially aimed for a larger number of perturbations per cell, but these have been challenging to achieve in our primary cells, and may require cells from CRISPRi^124^ engineered mice.

The large scale of our screen and the modular organization of the regulatory circuit opened a path to systematically tackle genetic interactions. First, our large-scale screen encompassed a relatively large number of cells with multiple perturbations per cell, but as this was a random sample, any particular combination was present in too few cells to directly estimate their effects. However, because of the organization of the regulators in co-functional modules, we could assess genetic interactions at the level of modules, testing for the prevalence of significant inter- or intra-module interactions globally, as well as their impacts on individual genes. This analysis showed that intra-module interactions are far more prevalent than inter-module interactions and impact specific target processes. Consistently, the impact of most inter-module combinations of perturbations can be quite well predicted by a naïve additive (linear) model. Moreover, learning the interaction effects was improved by comβVAE, a conditional variational autoencoder we developed that relies on the latent structure of the expression profiles. This shows the power of combining rich profiles and modular structures to allow prediction of unobserved experiments. Using a higher number of perturbations per cell, implementing ‘compressed screens’^123^,(Yao et al, bioRxiv 2023) and dedicated gene editing tools, such as Cas12^125^, should help further facilitate the dissection of genetic interactions.

### Leveraging cell screens to decipher human genetics

The reverse genetic approach of Perturb-Seq screens can complement and help interpret the results of forward genetic studies in humans, such as GWAS, especially in systems like primary DCs, where human cell models are lacking. By considering both regulators and regulated programs from our model in the context of associations from human GWAS and single cell profiles from relevant human disease tissues, we found that both common and low frequency variants in regulators in module M1 are enriched for heritability in immune-related traits, including the predicted E3 *WDR36* (in allergy/eczema and blood traits) and E3 adapter *PTPN11* in T1D and blood traits. Moreover, two of the programs affected by perturbations in module M1 regulators – ER stress (GP2) and translation (GP6) are differentially expressed in relevant cell types in immune disease, including macrophages and DCs in UC (GP2) and fibrosis and asthma (GP6).

### Limitations of our study

Although our study explores genetic circuits regulated by E3 family genes whose direct action is post-transcriptional, we rely on expression profiles as phenotypes, limiting our ability to draw direct mechanistic conclusions. We partly address this by *post hoc* analysis of protein-protein interaction and inference of TF activity, but further mechanistic studies and regulatory models that explicitly include physical interactions will be needed to generate a comprehensive model that is both causal (genetic) and mechanistic. For genetic interactions analysis, we were limited in our ability to transduce BMDCs at high MOI, resulting in fewer cells with multiple perturbations than desired. While module-level analysis allowed us to characterize genetic interactions and test our ability to predict them, this is a simplification. Moreover, our comβVAE currently focuses on inter-module interactions and further developments are needed to predict intra-module interactions, which are more prevalent. Finally, although our goal was to relate E3 family members and complexes in inflammatory circuits to human disease, no human DC line exists and patient-derived material is limited in scale and accessibility for genetic perturbations. We therefore screened mouse primary cells, and then related this signal to human genetics signal to prioritize regulators that may also play a large role in human health and disease.

## STAR Methods

### Mice

Six- to eleven-week-old female, constitutive Cas9-expressing mice were obtained from the Jackson labs (strain # 026179). All animal protocols were reviewed and approved by the MIT Committee on Animal Care (CAC protocol 0618-034-21) and all experiments conformed to the relevant regulatory standards.

### Bone marrow derived dendritic cells

BMDCs were differentiated and perturbed as previously described^16^. Cells were grown in RPMI media (ThermoFisher 21870-076) supplemented with 10% heat inactivated FBS (Invitrogen), 100 U/mL penicillin/streptomycin (GIBCO 15140122), 2 mM L-glutamine (ThermoFisher 25030081), 10 mM HEPES (GIBCO 15630080), 1 mM Na pyruvate (ThermoFisher 11360070), 1X MEM nonessential amino acids (VWR 45000-700), 55 μM β-mercaptoethanol (GIBCO 21985023), and 20 ng/mL recombinant murine GM-CSF (PeproTech 315-03). On day 0, bone marrow was extracted from mouse femur and tibia by cleaning surrounding tissue and crushing the bones gently via mortar and pestle. Bone marrow was filtered with a 70μm cell strainer, and red blood cells were lysed in 2 mL RBC lysis buffer (Sigma R7757) for 10 minutes at room temperature. RBC lysis was quenched with 18 mL media, cells were spun at 1,500 RPM and resuspended in 25 mL media. Following a final 70μm filtration, white blood cells were plated in 1,000mm non tissue culture-treated plastic dishes with 10 mL media at 200,000 cells/mL. On day 2, cells were fed with 10 mL media and infected with lentivirus (described below). On day 5, 12 mL media were removed carefully avoiding nonadherent cells, and 10 mL fresh media were added. On day 7, 5 mL media were added to cultures. On day 8, cells were collected, spun at 1,500 RPM and resuspended in 10 mL fresh media at 1x10^6^ cells/mL. On day 9, cells were stimulated for 3 hours with 100 ng/mL LPS (Invivogen, tlrl-peklps) and harvested by scraping. Cells then underwent antibody staining for cell hashing (described below) and mKate2^+^ (perturbation vector) and GFP^+^ (Cas9) cells were enriched by Fluorescence Activated Cell Sorting (FACS) (∼8% population) prior to single cell library generation (**Figure S1**).

### Selection of genes for perturbation

All 898 genes annotated as ‘E3 family’ were identified from the *Mus musculus* species in the iUUCD 2.0 database^7^ on April 2019. The ‘E3 family’ gene search included members with ‘E3 activity’, ‘E3 adaptor’ and ‘ULD/UBD’ designations in iUUCD. This list was supplemented with 1,054 *Mus musculus* genes identified by an NCBI Gene search of the term ‘E3 activity’, to a final non-redundant list of 1,137 E3s and interaction partner genes.

### Design and construction of feature barcoding lentiviral vector

To generate a lentiviral perturbation vector compatible with Perturb-seq Feature Barcoding technology (10x Genomics), a new FB-LentiGuide-Puro-mKate2 vector was designed (pRDA_122: Supplemental Sequence File; AddGene #TBD), where sgRNAs contain a 3’ terminal binding sequence (gctcacctattagcggctaagg) complementary to Feature Capture oligos present on 10X v3 and v3.1 beads. To enable FACS or puromycin selection of perturbed cells, expression of mKate2-2A-PuroR was driven by the Ef1α promoter. The designed vector was generated by GenScript.

### Cloning of guide pools

A 3,390 Perturb-Seq guide library was designed with three guides targeting each of the 1,130 genes using the Broad Institute Genetic Perturbation Platform Web sgRNA Designer (https://portals.broadinstitute.org/gpp/public/analysis-tools/sgrna-design)^126^, along with 330 control guides (165 nontargeting guides and 165 targeting intergenic regions). The pooled CRISPR library was cloned into the FB-LentiGuide-Puro-mKate2 vector backbone as previously described^17^.

### Cell hashing and overloading

BMDCs were washed with PBS and resuspended in 50μL PBS+ 2% BSA and 0.1% Tween20 and stained with BioLegend hashing antibodies (BioLegend 155801, 155803, 155805, 155807, 155809, 155811, 155813, 155815) as previously described^127^ and mKate2^+^GFP^+^ cells were sorted by FACS (**Figure S1A**). 40,000 cells were loaded per channel on 46 3’ v3 channels according to the manufacturer’s instructions^128^.

### Single cell RNA-seq library generation

ScRNA-seq libraries were generated following the manufacturer’s instructions (v3 User Guide, 10x User Guide, with Feature Barcoding)^128^.

### Feature barcoding (gRNA) and hashtag libraries generation

The cDNA amplification reaction was mixed by adding 5 μL of 2 μM HTO additive (**Table S14**), 15 μL Feature cDNA primer 2000096, 50 μL Amp Mix (10x Genomics, 2000047 or 2000103) and 30 μL cDNA followed by PCR (98°C for 3 min; 98°C for 15 sec, 63°C for 20 sec, 72°C for 60 sec x 11 cycles; 72°C for 1 min; hold at 4°C). cDNA supernatant was selected by collecting the eluant from the 0.6X SPRI cleanup of the cDNA amplification reaction. Eluant was taken from the 0.6X SPRI cleanup of the cDNA amplification reaction, and another 60μL of SPRI beads were added to the 150μL cDNA supernatant. After performing two 80% ethanol washes, elution was performed in 50μL EB buffer (https://www.qiagen.com/us/products/discovery-and-translational-research/lab-essentials/buffers-reagents/buffer-eb/). An additional 1.0X SPRI elution was performed in 30μL EB buffer. This SPRI-purified cDNA supernatant was used as template for both hashtag and feature barcoding library generation.

To generate feature barcode (single cell gRNA) libraries, 50μL 10X Amp mix was next mixed with 45μL Feature SI Primers 1 and 5μL SPRI-purified cDNA supernatant followed by PCR (98°C for 45 sec; 98°C for 20 sec, 60°C for 30 sec, 72°C for 20 sec x 15 cycles; 72°C for 1 min; hold at 4°C). Product was purified with 1.0X SPRI beads, eluting in 30μL EB. Next, the product was run on a 2% TBE agarose gel at 140 volts for 40 minutes and the 250-300 bp gel fragment was purified by adding 800μL agarose dissolving buffer (https://www.zymoresearch.com/products/adb-agarose-dissolving-buffer) to each sample and incubating at 55°C at 1,250 RPM for 10 minutes. The dissolved gel was added to a Zymo DNA Clean & Concentrator-5 kit column and columns were spun for 30 seconds at maximum speed, followed by two 200 μL washes with the included wash buffer (https://www.zymoresearch.com/products/dna-clean-concentrator-5). Columns were spun for four minutes at max speed to dry, then transferred to a new tube followed by elution in 15μL water (D4014). 5μL gel purified product was mixed with 50μL 10X Amp mix, 35μL Feature SI Primers 2, and 10μL Chromium i7 sample indices (PN-120262/PN-220103: Chromium i7 Multiplex Kit) followed by PCR (98°C for 45 sec; 98°C for 20 sec, 54°C for 30 sec, 72°C for 20 sec x 6 cycles; 72°C for 1 min; hold at 4°C) and the product was purified with 0.7X SPRI and eluted in 30μL EB.

To generate hashtag libraries, 5μL SPRI-purified cDNA supernatant was next combined with 25μL NEBNext 2X MasterMix (https://www.neb.com/products/m0541-nebnext-high-fidelity-2x-pcr-master-mix#Product%20Information) and mixed with 19μL water, 0.5μL 10μM SI-PCR primer, 0.5μL 10 μM K_HTOX primer (**Table S14**) followed by PCR (98°C for 10 sec; 98°C for 2 sec, 72°C for 15 sec x 21 cycles; 72°C for 1 min; hold at 4°C). Product was purified with 2.0X SPRI and eluted in 15μL TE buffer (https://www.thermofisher.com/order/catalog/product/AM9849).

### Library sequencing

Gene expression, hashtag, and feature barcoding libraries were pooled and sequenced together across 46 Illumina HiSeq 2500 lanes (R1 28 I1 8 R2 90), yielding on average 15,900 scRNA-Seq reads per cell (R3: 350M reads per lane total of 12 lanes, 78% spike-in), 680 hashing reads per cell (R3: 15M reads, 4% spike-in), and 3,600 feature barcoding reads/cell (R3: 80M reads, 18% spike-in).

### Read alignment and demultiplexing

ScRNA-seq reads were mapped to the reference mouse genome mm10_v3.0.0 with Cumulus^129^ executed through Terra (https://app.terra.bio/), using CellRanger 3.0.2. Demultiplexing of cell-hashing and feature barcoding data was performed with DemuxEM^34^ as implemented in Cumulus.

### scRNA-seq preprocessing

Empty droplets were removed at FDR > 0.01 by EmptyDrops ^36^ with parameters: lower= 200, niters= 10,000, ignore=10 and retain= 1000. Droplets with either <300 detected genes, <1,000 total UMIs or >15% mitochondrial reads were removed, retaining 1,071,671 droplet profiles. 13,811 genes expressed in at least 400 of these droplets were retained. 838,201 droplets were categorized as singlets based on hashtags by DemuxEM^34^. Cells were assigned a feature (guide) if they had at least 2 feature barcode UMIs, and feature barcode-UMI pairs with <20% of the reads per cell were removed^130^, yielding 341,664 cells assigned one barcode and 177,871 cells assigned at least two. 186 targeting guides detected in <20 single perturbed cells were removed from further analysis, retaining 3,204 targeting guides.

### scRNA-seq expression matrix and dimensionality reduction

Single cell expression matrix and feature barcodes were processed in an anndata object format in Scanpy^131^. Raw counts were saved in the ‘counts’ layer for downstream analysis. Expression counts per cell were normalized, to a total of 10^4^ counts per cell, and normalized values were log transformed (natural log), after adding a pseudocount of 1.

A *k*-nearest neighbor (*k*=15) cell neighborhood graph was constructed with the first 50 principle components (PCs) of the log normalized expression matrix and clustered with the Leiden algorithm^132^ (resolution=0.5). Gene signatures were computed as the average expression of the gene set in the cells minus the average expression of a reference set of genes that is randomly sampled from the same expression bins.

Outlier control guides were identified by PCA of the log-normalized expression matrix of the 44,074 control cells with one of 330 control guides, followed by fitting a linear regression model to each of the top 100 PCs with Python statsmodels package^133^, where in each model one PC was the response and the binary feature barcode matrix of the control guides were the covariates. To identify outliers, the 330 X 100 coefficients matrix was fitted with each of four algorithms in scikit-learn Python package^134^: isolation forest^135^, elliptic envelope^136^, local outlier factor^137^, and one-class SVM, and the 9 non-targeting and 22 intergenic guides that were predicted as outliers by at least three methods were removed.

### Prediction of corresponding gene expression clusters in unperturbed cells with and without LPS stimulation

Cluster assignment of LPS unstimulated unperturbed and LPS stimulated unperturbed cells were predicted by a logistic regression model trained on the LPS stimulated perturbed dataset, with the 10 cluster scores of the top 100 marker genes of the clusters as covariates.

### Guide and knockout enrichment analysis

To test guide depletion in the screen *vs*. the initial guide pool, distributions of the ratio between the number of cells assigned a (single) guide in the screen *vs*. number of guides in the initial pool were generated for each of the 3,390 targeting and 330 control guides. The ratio distribution of the control guides was taken as background, to calculate an empirical P-value of the depletion of each targeting guide. Targeting guides with ratio of at most 0.08587 were identified as depleted (CDF_null_ (0.08587) = 0.0498).

One-sided Fischer’s exact tests were used to test the enrichment (separately, depletion) of cells with a particular guide (or guides targeting the same gene) in each cell subset, where the test schema was the tested group versus rest, and a Benjamini-Hochberg FDR was calculated.

### Identification of congruent guides targeting the same gene

The effects of each of 3,204 targeting guides on 6,685 genes expressed in at least 5% of the 341,664 singly-perturbed cells were learned using a negative binomial regression model, with control cells as reference, and correcting for the total number of detected genes in a cell, % mitochondrial reads and cell states identified by the initial Leiden clustering. The pairwise Pearson correlation coefficient between the effect size profiles of each pair of guides targeting the same gene were calculated. If no pairs had a positive correlation, all guides were discarded. If all three pairs had *r*> 0.015, all three guides were retained. Otherwise, only the guide pair with the highest positive correlation was retained.

### A linear regulatory model of knockout effect

A mixed effects negative binomial linear regression model was fit for each of the 6,685 affected (response) genes, where the gene expression values were the response variable, the cell states identified by Leiden clustering were the random effect covariate, and the knockout target gene confounders (number of detected gene/cell, %mitochondrial reads/cell) were the fixed effect covariates, and control cells were the reference. Benjamini-Hochberg FDR was used to correct for multiple hypotheses (6,658 tested genes) with a stringent threshold, such that most regulatory coefficients close to zero were not significant. To generate a background distribution for the number of genes significantly affected due to lentivirus infection or off-target effects, the same model was fit for each of the 299 control guides, testing one control guide against the rest of the control guides. Based on this background distribution, 329 perturbed genes were retained as ones with significant (FDR < 0.1) effect on at least 15 of the 6,685 tested genes.

### Identification of co-functional modules and co-regulated gene programs

The 329 retained knockouts (perturbed genes) were grouped based on their effects on the 1,041 genes that were affected by at least by 4 perturbations. To this end, the top 50 PCs of the scaled and centered effect size matrix B (B_ij_ = effect size of knockout of gene *i* on gene *j*) were used to calculate a *k*-nearest neighbors graph (*k*=10) of the knockout (perturbed) genes, and the Leiden algorithm (resolution=0.64) was used to identify the 6 clusters as co-functional modules.

Similarly, to identify co-regulated gene programs, the top 50 PCs of the scaled and centered B^T^ were used to construct a *k*-NN graph of the 1,041 response genes (*k*=10), and the Leiden algorithm (resolution=0.8) was used to identify 8 gene programs. Three of these programs were selected for further subclustering upon manual inspection, resulting in 11 gene programs.

### Embedding cells jointly on KO module information and their gene expression profiles

To assess the change in cell distributions across DC2.1, DC2.1 and DC2.3 subsets upon perturbations, supervised-UMAP ^138^ was used (target_weight = 0.5, kNN n_neighbors=18) to embed cells based on their normalized expression profiles and module assignment (M1-M6).

### Calculating Wasserstein distances within and across modules

The average population distances between cell subsets perturbed for members of each module (or controls) or between randomly sampled cell subsets perturbed for members of the same modules, were calculated by sampling without replacement 300 cells (100 times), computing Wasserstein distances between pairs of cell populations using the Python Optimal Transport Library ^139^, and averaging across 100 iterations.

### Protein-protein interaction analysis

*Mus musculus* protein-protein interaction network data was downloaded from STRING DB (version 11.5) and interactions with experimental evidence score > 0 were selected. Interactions between the 329 knockout targets were used to generate a protein-protein interaction graph. To test for enrichment of intra- and inter-module interactions, 400 random degree-preserving graphs were generated using the BiRewire R package^140^ and the distribution of number of intra- and inter-module interactions in these graphs was used as the null distribution to calculate empirical P-values for the corresponding observed number of interactions.

### Inference of KO effects on TF factor activity

Expression scores of high confidence targets (Levels A and B) activated by 123 mouse TFs in DoRothEA^97, 141, 142^, were calculated using the R package decoupleR^96^. A linear regression model was used to infer the effects of each 329 KOs on the expression of targets of each of the 123 TFs, where in each model the response variable was the expression score of each TF’s target genes, and the covariates were the 1-hot encoded feature barcode matrix (with control cells as the base level) and possible confounders (cell clusters, %mitochondrial reads, number UMIs).

### ICA module factorization

Independent components analysis^99, 143^ was used to identify statistically independent factors from the 1041 X 329 effect size matrix β from the mixed effect linear model (*b_ij_* = estimated effect of knockout of gene *j* on expression of gene *i*). A source of variation *S* = {*S*_1041*k*_, *S*_2*k*_, *S*_3*k*_, … *S*_1*k*_ } is defined as the set of relative weights (*i.e.*, relative expression states) of genes 1, … , 1,041, such that the effects of perturbation *j* on expression of gene *i* is a weighted sum of the effects over *P* different sources, written as:

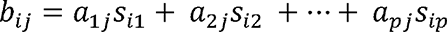

where *a*_1*j*_, *a*2_*j*_, …, *a*_*pj*_ are the mixing weights and *a*_*pj*_ is the overall effect of perturbation *j* on source *p*. Thus, in matrix β, each row is an observation of a gene’s expression changes due to the varying effects of the knockouts on various pathways (sources of variation) to which the gene belongs.

Identifying the *P* underlying pathways *s_1_*,…,*s_P_*affected by 329 knockouts is formulated as finding a source matrix *S* (1,041 by *P*) and mixing matrix *M* (*P* by 329) which are both unknown:

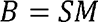

ICA relaxes this factorization problem by assuming that the source signals are independent and requiring that they be non-Gaussian ^99^. Although modeling total perturbation effects as a linear combination of factors may miss nonlinear relationships, the nonlinear separation problem is not identifiable.

ICA decomposition was computed using Information-Maximization (Infomax)^144^, as implemented in the ICA package in R^145^. The optimal number of latent sources, *P* was determined considering (**1**) the number of non-Gaussian components estimated by the Ladle estimator^146, 147^; (**2**) reconstruction error of the original matrix from the obtained statistically independent components; and (**3**) prediction power of the identified factors for the effects of unseen perturbations during fitting. For (**2**), the ICtest R package^146^ was used to compute the ladle estimates (gn) for different number of factors, where ‘gn’ is the sum of the vectors giving the measures of variation of the eigenvectors and the normalized eigenvalues of the fourth order blind identification (FOBI) matrix and the estimated number of Gon-gaussian components is the value where gn takes its minimum. For (**3**), for *P* between 2 and 30, 80% (263) of the 329 perturbations were randomly sampled 10 times, ICA was fitted each time to the 1,041 by 263 matrix and the effects of each of the remaining 66 perturbations was predicted with a simple linear regression model, where the 1,041 by P matrix *S* was the covariate matrix.

After selecting *P*=15, the full 1,041 X 329 effect size matrix β was decomposed and for each factor in *S,* gene *i* was defined as a prominent gene defining this factor if it had outlier weights *s*_*ik*_ < *Q*_1_(*s*_*k*_) - 1.5(*Q*_3_(*s*_*k*_) - *Q*_1_(*s*_*k*_ | *s*_*ik*_ < *Q*_3_(*s*_*k*_) - 1.5(*Q*_3_(*s*_*k*_) - *Q*_1_(*s*_*k*_. Likewise, a KO of gene *j* was defined as highly affecting factor *k* if it had outlier weights in component of the mixing matrix M, *m*_*jk*_ < *Q*_1_(*m*_*k*_ - (*Q*_3_(*m*_*k*_ - *Q*_1_(*m*_*k*_)) | *m*_*jk*_ > *Q*_3_(*m*_*k*_ - (*Q*_3_(*m*_*k*_ - *Q*_1_(*m*_*k*_)).

### Genetic interaction analysis

For inter-module interactions, for each of the 1,041 response genes, linear regression models were fit as follows:

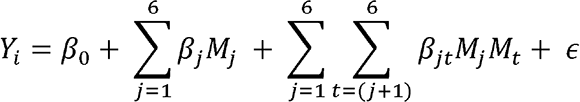

where *Y*_i_ is the normalized expression level of gene *i* (corrected for cell cluster, % mitochondrial reads and number UMIs), is a binary covariate denoting if the cell had a perturbation in a gene from module *i*. The model was fit with single KO cells and inter-module double KO cells. P-values of the β estimates were corrected with Benjamini-Hochberg FDR for the 1,041 tested genes.

For intra-module interactions, for each KO module we randomly partitoned (for 50 times) the KO module *M_j_* into two equal bins in terms of the number of KO genes, *M*_*j*_1_ and *M*_*j*_2_, and for each response gene *i* fit the model:

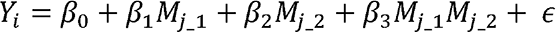

where *Y*_*i*_is the normalized expression level of gene *i* corrected for confounders as above. Benjamini-Hochberg FDR was first calculated for the p-values of the estimates for each of the 50 iterations, β estimates for which FDR >= 0.1 were set to zero, and then the mean of the 50 parameter estimates as taken as the inferred intra-module interaction effect.

### Deep learning model to predict interactions

To learn models that predict the effect of combinatorial perturbations, conditional VAEs (CVAEs) ^101^ were used, which model the distribution of a high-dimensional output as a generative model conditioned on the auxiliary covariates. In general, CVAEs aim to learn the marginal likelihood of the data in such a generative process:

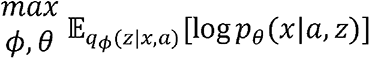

where *x* ϵ ℝ^*D*^ is a dataset of samples with labels (conditioned variable) *a* and generated by ground truth factors *z*, *ϕθ* while parametrize the distributions of the CVAE encoder and the decoder respectively. This can be rewritten as:

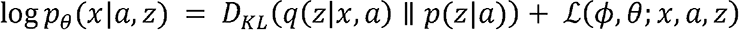

where *D*_*KL*_(||) is the non-negative Kullback-Leibler (KL) divergence between the true and the approximate posterior and ℒ(*ϕ, θ; x, a, z*) is the evidence lower bound (ELBO) on the log-likelihood of the data:

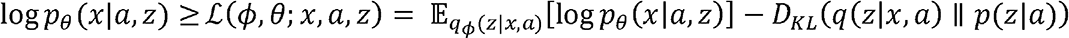

To make optimization tractable in practice, is typically set to the isotropic unit Gaussian 𝒩 (0,*I*)^148^. The ELBO for the VAE and CVAE factorizes across the samples^101, 148, 149^. Therefore, it is straightforward to apply computationally efficient minibatch based stochastic gradient descent (SGD) and learn the parameters *ϕ* of the encoder (*q*_ϕ_(*z*|*x,a*) and θ of the decoder by deep neural networks.^148^

In our com VAE model, (*x_i_*,*a_i_*) denotes a single data point *i*, where *x* ϵ ℝ^*G*^ is the G-dimensional observed log-normalized gene expression profiles of *G* genes in a single cell *i* and representing the *P*-dimensional auxiliary (independent) discrete covariates of the same cell, such as the knockout(s) perturbing the cell, or confounders (cell subtype or cell cycle phase). The perturbation covariates in are assumed to be independent binary covariates and a cell can have multiple perturbations, while other covariates are one-hot encoded. The latent variables z*_i_* are genenrated conditionally on a *D* dimensional vector *v* ϵ ℝ^*D*^ which are the embeddings of learnt with a single hidden layer neural network of D units which is jointly trained with the encoder-decoder framework. An adjustable hyperparameter β is introduced to the original CVAE objective, which was previously shown to result in more disentangled latent representations z in standard VAE models ^150, 151^ :

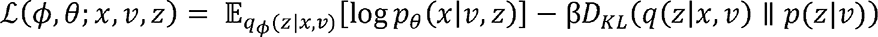

Assuming the data generating process described above, our objective is to train a model such that, a target counterfactual distribution of gene expression *X*_*i*_^′^ can be generated if cell *i* had the covariates instead of . For the network architecture after benchmarking the number of hidden units [2,3,4,5,6], and number of units per layer [32, 64, 128, 512, 1024], the encoder and decoder networks were defined with 2 hidden layers, with 512 units in each layer, and ReLU (rectified linear unit) activation function used between the hidden layers. The dimensions [16,32,64,128] were benchmarked for the dimensions of *z* and and both were set to 64. Models were trained and benchmarked with log-transformed normalized gene expression values of 1,041 genes corrected for cell clusters, % mitochondrial reads and total UMI counts to minimize the effect of confounders in evaluating generated *vs*. observed effects. The model was implemented in Pytorch, and trained with hyperparameters batch_size =1000, max_epochs=8000, optimizer=adam, learning_rate= 0.001, weight_decay=0.

### Disease progression gene programs

Disease progression programs were defined as previously described^102^ using publicly available scRNA-seq datasets, processed, annotated, and analyzed as previously described^102^. A gene-level non-parametric Wilcoxon rank sum differential expression test was performed between cells from healthy and disease tissues of the same cell type as previously described^102^.

### Identification of heritability signal

Gene programs and co-functional modules were tested for enrichment in heritability signal using both scLinker^102^ and MAGMA^103^ over a set of 60 relatively independent diseases and traits (average N = 297K)^102^.

In sclinker, each program or module were combined with enhancer-gene linking strategies defined by either SNPs in enhancers linked to genes based on the Roadmap^152^ and Activity-By-Contact (ABC) SNP-To-Gene (S2G) strategies either aggregated across all biosamples related to blood (RoadmapUABC-Blood). For each gene score X and S2G strategy Y, a combined annotation X × Y was defined by assigning to each SNP the maximum gene score among genes linked to that SNP (or 0 for SNPs with no linked genes); this generalizes the standard approach of constructing annotations from gene scores using window-based strategies^153, 154^. Heritability analysis of these sclinker annotations was performed using stratified LD score regression^155, 156^ conditional on a set of 86 baseline coding, conserved and LD-related annotations (baseline- LDv2.1^157^). The Enrichment Score (Escore) metric^102^ reported was derived from heritability enrichment analysis and its corresponding p-values.

For MAGMA analysis, the MAGMA z-score was computed for each gene module or program using a 0kb window based strategy for linking SNPs to genes^103^ and then a gene set enrichment analysis of the MAGMA z-scores was performed for each with respect to a set of 1,000 sets of same size of randomly selected genes from across all perturbation programs (using the fgsea software^158^).

## Acknowledgements

We thank L. Gaffney and A Hupalowska for assistance with figure design and graphic, and Brian Cleary, Romain Lopez and members of the Regev and Theis labs for fruitful discussions. Work was supported by the Klarman Cell Observatory, National Human Genome Research Institute (NHGRI) Center for Excellent in Genome Science (5RM1HGP706193-09), and HHMI (AR), and NIH F32 Ruth L. Kirschstein Postdoctoral Fellowship (5F32AI138458) (PT).

## Author Contributions

Conceptualization: KGS, AR Perturb-seq vector design: JD, KGS

Expanded Perturb-seq design and experimentation: KGS, OK, PIT, LZG, TD, DP, RR, NS Sequencing: KGS, TD

Computational analysis design and execution: BE, KGS, KD, KJ, KDY Supervision: AR, ORR

Writing – original draft: KGS, BE, AR, KD, KJ Writing – review and editing: all authors

## Data and materials availability

scRNA-seq data generated in this work are available from the Gene Expression Omnibus with accession number TBD and the Single Cell Portal with accession number TBD. Code used in analysis of sequencing data is available from GitHub link https://github.com/EraslanBas/PerturbDecode.

## Supplementary Tables

Table S1. Curated E3 gene family list

Table S2. Perturb-seq guides and oligo pools

Table S3. Gene lists for cell type signatures

Table S4. Top 1000 differentially expressed genes in cell clusters

Table S5. Enrichment or depletion of perturbed genes compared to the input library

Table S6. Enrichment or depletion of perturbed genes in cell subsets and clusters

Table S7. Regulatory coefficients in the model

Table S8. Gene programs and regulatory modules

Table S9. E3 and E3 complex member regulator metadata

Table S10. ICA factor genes, regulators and explained variance

Table S11. Regulatory coefficients in a model with genetic interactions

Table S12. Immunologic trait heritability in modules

Table S13. Extended sc-linker program heritability

Table S14. Primers for sequencing library generation

## Supplementary Materials

**Figure S1, related to Figure 1.**
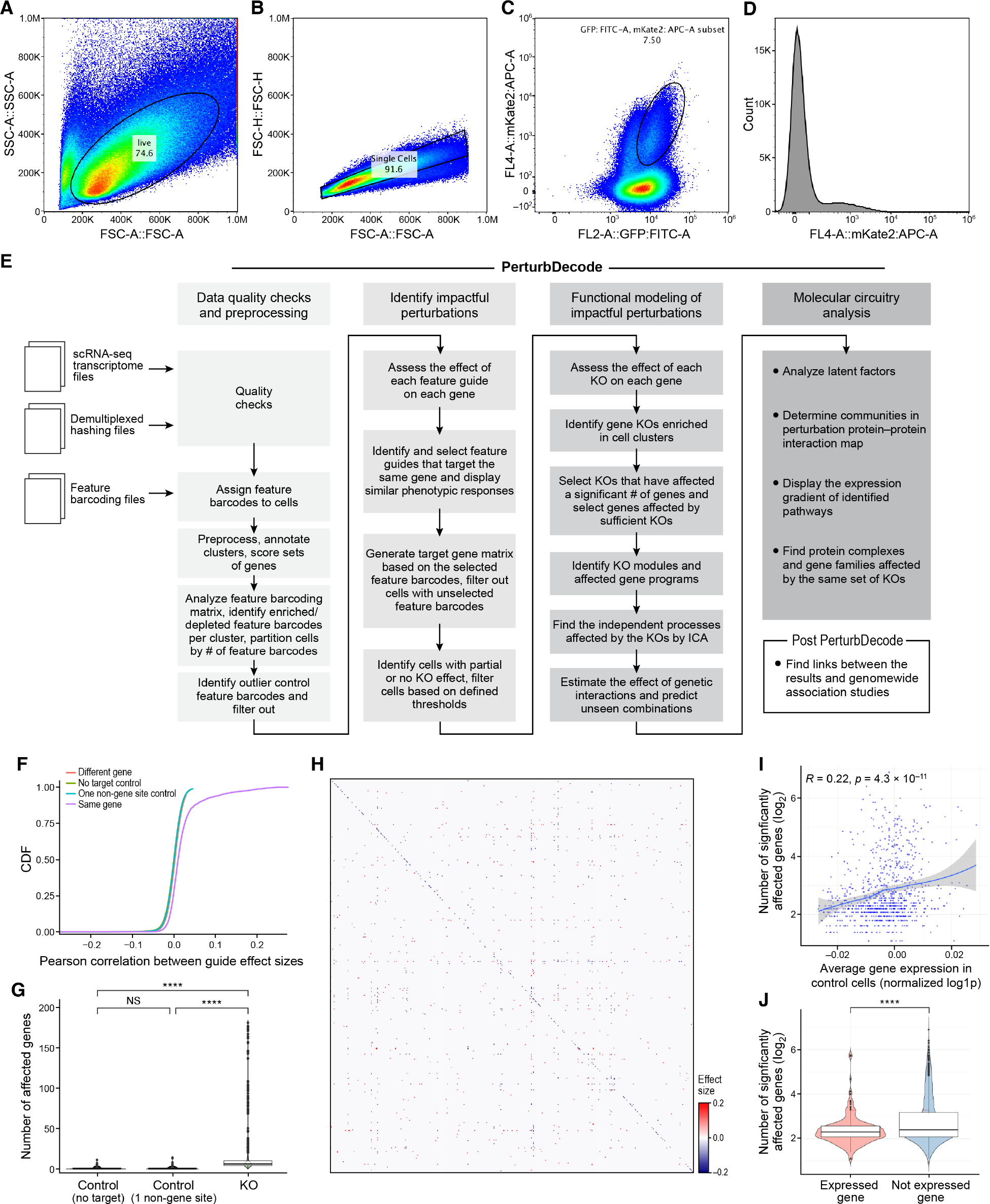
Design and performance of large-scale Perturb-Seq in primary BMDCs **A-D.** Sorting of perturbed cells for profiling. **A,B.** Forward scatter (x axis) versus side scatter (y axis) for selected live (A) single (B) perturbed BMDCs. **C.** GFP fluorescence (x axis; Cas9 mice cells) versus mKate2 fluorescence (y axis; Perturb-Seq vector) to select mKate2^+^GFP^+^ cells. **D.** Distribution of mKate2 expression (x axis) in sorted live, single cells. **E.** PerturbDecode. Detailed workflow. **F-J.** Quality control for impact of guide-based perturbations. **F.** Cumulative distribution function (CDF) (y-axis) of Pearson’s *r* (x-axis) between the effect sizes of guides targeting the same gene (purple), different genes (red), one gene and one no-target control (green), or one gene and one intergenic control (blue). **G.** Distribution of number of genes (y-axis) (of 6,685 tested genes) significantly affected (FDR < 0.1) by non-targeting controls, intergenic controls or targeting guides (with guides targeting the same gene combined) (x-axis) (**STAR Methods**). **** P < 2.2*10^-16^, one-sided Wilcoxon rank-sum test. **H.** Significant effect sizes (color bar; blue/red negative/positive fold-change; FDR<0.1) of perturbing each of 544 targets (rows) that were also among the 6,685 genes with tested expression on itself and the other 544 targets (columns). Rows and columns are ordered alphabetically. 137 of 539 genes significantly negatively affected their own expression (diagonal). **I.** Number of genes (y axis) whose expression is significantly (FDR<0.1) affected by perturbation of each of 849 perturbed genes (from 13,811 detected genes) and the mean expression of these perturbed genes (x axis, normalized log1p). Pearson’s *r* and significance in the upper right corner. **J.** Distribution of number of genes (y axis) significantly affected (FDR < 0.1) by the perturbation of genes that are (“expressed”) or are not (“not expressed”) in the 13,811 detected genes. **** P-value < 10^-4^, one-sided Wilcoxon rank-sum test.

**Figure S2, related to Figure 1.**
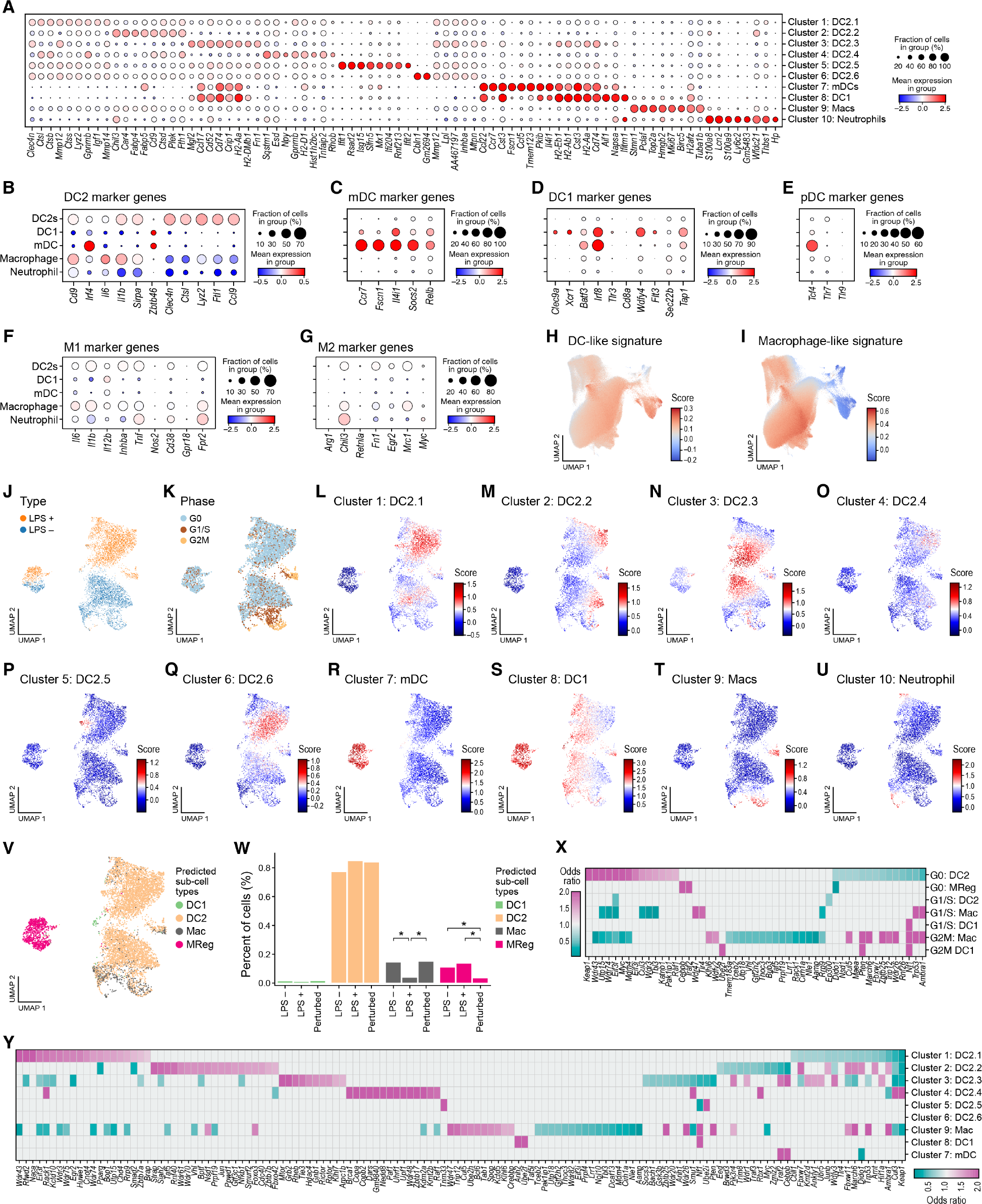
Proportions of cells in different states and subtypes in screen are affected by perturbations. **A-G**. Markers of specific cell subsets in the screened BMDC population. Mean expression (dot color, mean normalized log1p expression) and fraction of expressing cells (dot size) for genes differentially expressed (columns) in each of the 10 cell clusters (A, rows) or in the major cell sub-types (B-G, rows). **H,I.** DC- vs. macrophage-like gradient. UMAP embedding of 519,535 cell profiles (as in Figure 1C) colored by DC (DC1+DC2+mDC genes) or macrophage signature score. **J-V.** Cell subsets and stated in unperturbed resting or LPS-stimulated BMDCs. UMAP embedding of 3,655 unperturbed unstimulated and 4,027 unperturbed and LPS stimulated (3h) BMDC profiles colored by treatment (J), inferred cell cycle phase (K), signature scores for the top 100 upregulated genes of each of the 10 clusters of Figure 1C (**L-U**), or their predicted major cell subtype (**V**) (**STAR Methods**). **W.** Perturbation and stimulation affect cell subtype proportions. Percentage of cells (y axis) of each of four major subtypes in the screen (color legend) in unperturbed unstimulated, unperturbed LPS stimulated and perturbed, LPS stimulated data (x-axis). * P < 2.2*10^-16^, one-sided Fisher’s exact test. **X,Y.** Specific perturbations are enriched or depleted in specific cell subsets and clusters. Odd-ratio (color bar) of enrichment (pink) or depletion (blue) (FDR < 0.15, one-sided Fisher’s exact test) of cells with a perturbed gene (rows) in cell cycle phases in major subtypes (X, columns) or in the 10 cell clusters (Y, clusters as in Figure 1C).

**Figure S3, related to Figure 2.**
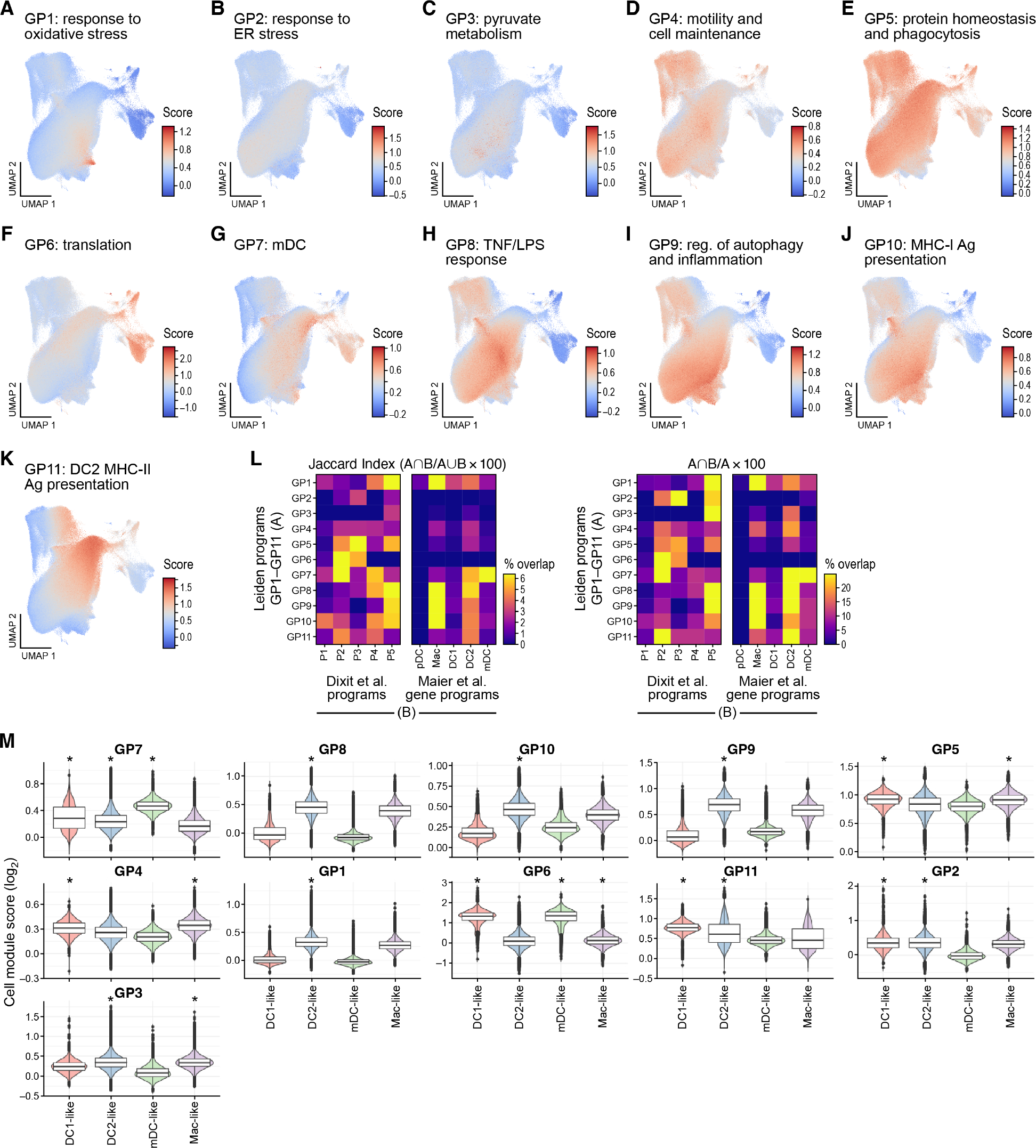
Gene programs in the regulatory model **A-K.** Gene programs expressed in different cell subsets. UMAP embedding of cell profiles (as in Figure 1C) colored by expression scores of each program genes. **L,M.** Relation of gene programs to other programs and cell subset signatures of BMDCs. **L.** Jaccard index (left) and fractional overlap (right) between each gene program (rows, “A”) and programs in an earlier small Perturb-Seq screen of 24 TFs in the LPS-stimulated BMDCs ^16^ (left, columns, “B”) or DC subset signatures ^39^ (right, columns, “B”). **M.** Distribution of program scores (y axis) for DC1-, DC2-, mDC-, and macrophage-like cell subsets (x axis). * P < 0.05 one-*vs.*-rest one-sided Wilcoxon rank sum test.

**Figure S4, related to Figure 3.**
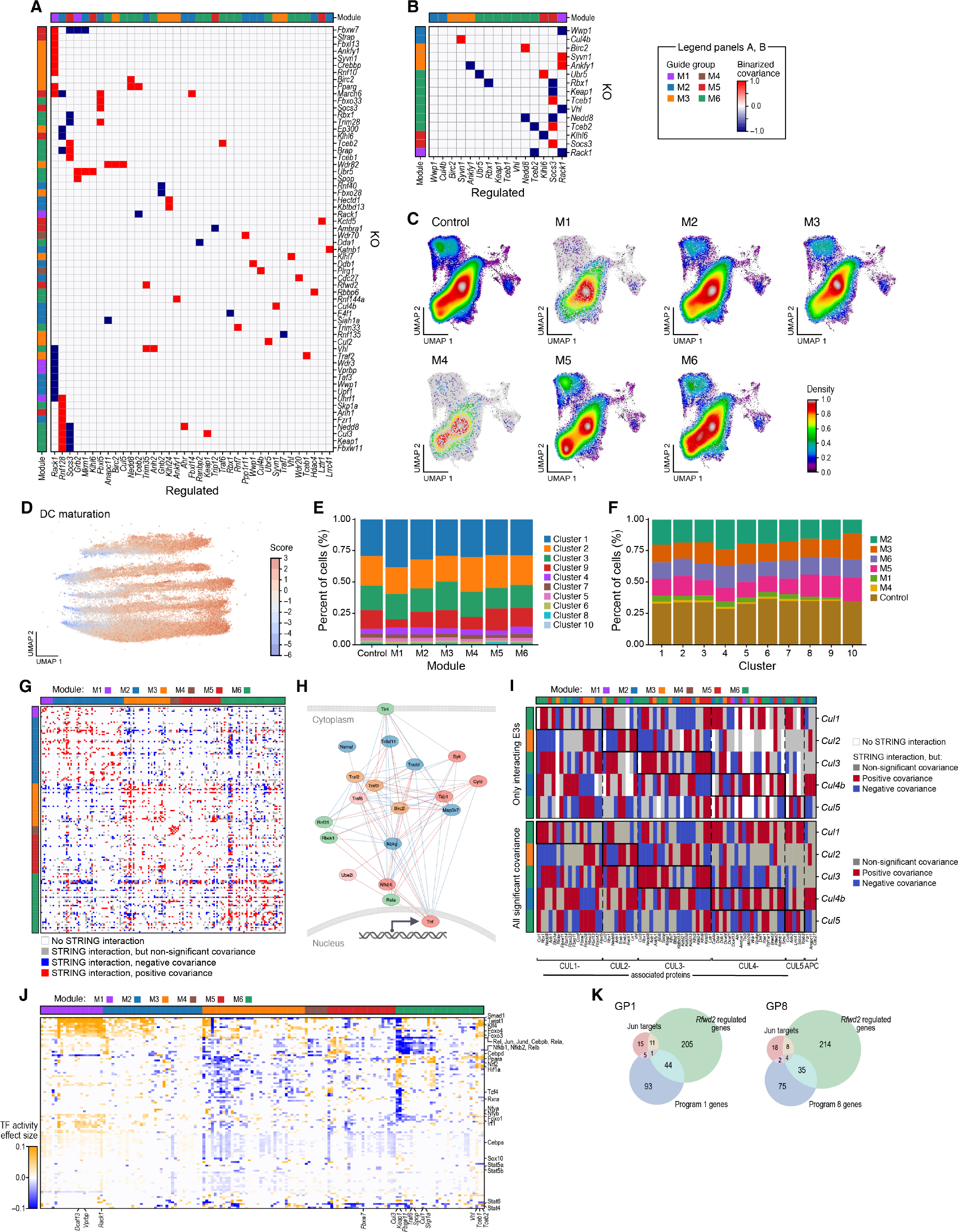
Relation between regulatory (genetic) and physical (protein-protein and TF-target) interactions. **A,B.** Three E3 ligases are ‘authorities’ highly regulated by other E3s. Binarized regulatory effects (blue/red: negative/positive) of perturbing each of 60 E3 ligases in the model that significantly affected the expression of at least one other of the 60 E3 ligases, for all 60 E3 ligases (A) or for the 15 that both impact and are impacted by another E3 (B). Module membership is labeled by color on left and top. Negative effects of the KOs on its own RNA level are not shown in A and shown in B. **C-F.** Co-functional modules affect distribution of cell states. **C.** UMAP embedding of cell profiles (**as in** Figure 1C) colored by Gaussian kernel density estimations of cells with control guides (top left) or guides targeting genes with each module (label on top). **D.** Supervised UMAP embedding of DC2.1, DC2.2, and DC2.3 cell profiles using a cells’ co-functional module assignment as the response label (as in Figure 3C, **STAR Methods**), colored by the difference of macrophage and DC signature Z scores. **E.** Percentage of cells (y axis) with guides targeting genes in each module (x axis) belonging to each of the cell clusters of Figure 1C (colors). **F.** Percentage of cells (y axis) in each of the cell clusters of Figure 1C (x axis) with guides targeting genes in each module (colors). **G-I.** Congruence between protein interactions and genetic effects. **G.** Physical interactions (blue, red, grey; experimental score > 0, STRING DB) or lack thereof (white) between each pair of 165 E3 ligases among the 329 regulators (rows, columns). Red/blue: the regulatory profiles of the physically interacting genes have significant (P<0.05) positive/negative correlation. Grey: the regulatory profiles of the physically interacting genes are not significantly correlated. Genes are sorted by module membership (colors on top and left). **H.** Physical interactions (edges, STRING DB experimental score > 0) between NFkB signaling pathway components included in the regulatory model (nodes), colored by significant (P<0.05) positive (red) or negative (blue) correlation of their perturbation effects. **I.** Top: physical interactions (blue, red, grey; experimental score > 0, STRING DB; color code as in G) or lack thereof (white) between each perturbed CLR E3 ligases (rows) and their CLR complex members, including adaptor domain proteins (columns). Columns are ordered by CLR physical complexes / interactions (boxes and dashed lines). Bottom: As on top, except all significant covariances are displayed regardless of interaction evidence. **J,K.** TFs impacted by and mediating the effects of E3 perturbation. **J.** Inferred activity scores (colorbar) of 109 TFs (columns) whose target genes are significantly (FDR < 0.1) induced (yellow) or repressed (blue) (by at least 10 perturbations) when perturbing each of 156 E3 and related genes (rows) (that affected at least 10 of the 109 TFs) (**STAR Methods**). **K.** Intersection between TF targets (DoRothEA), E3 expression targets, and gene programs.

**Figure S5, related to Figure 4.**
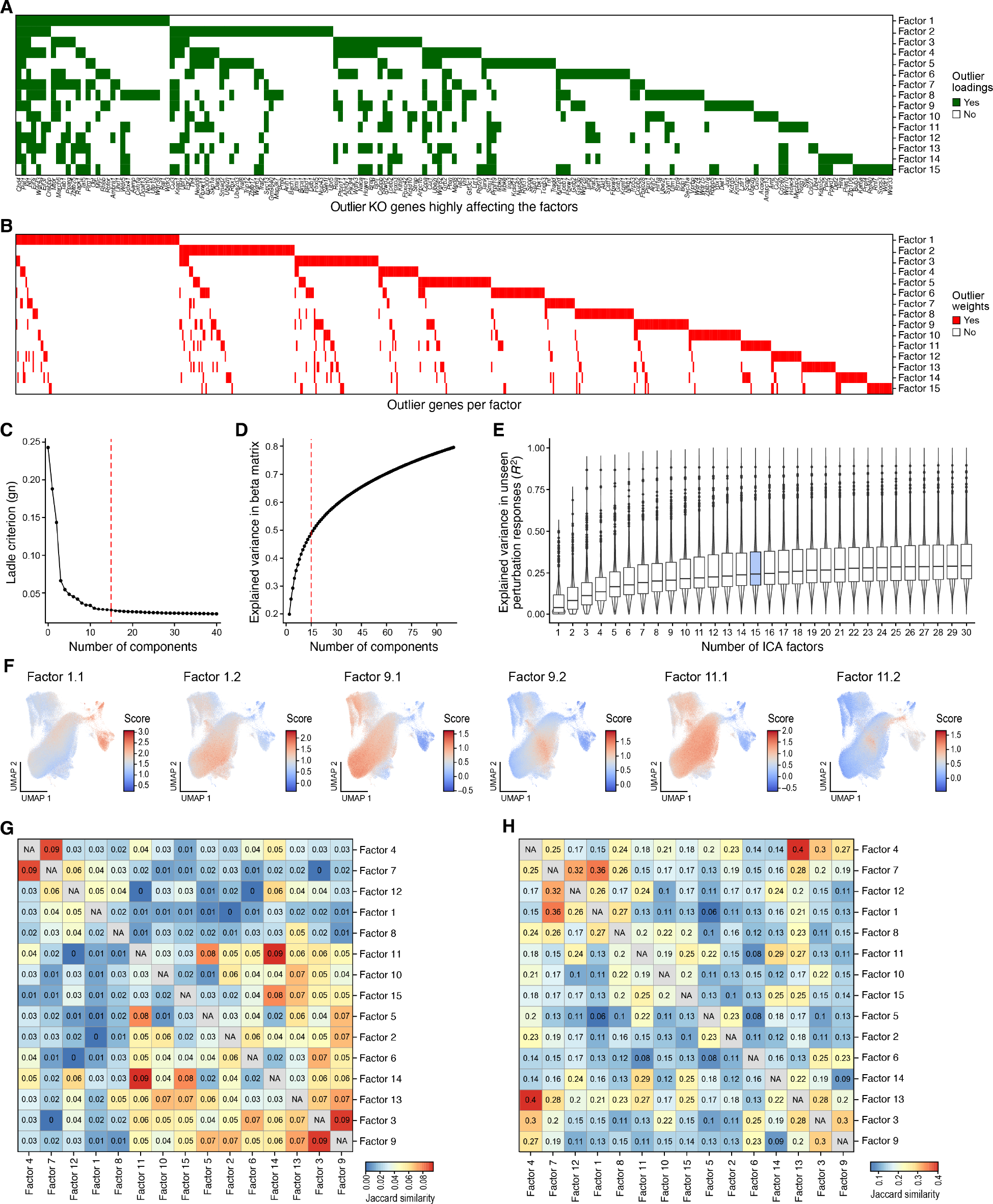
Independent factors in the regulatory model **A,B.** Genes and regulators associated with each factor. Regulators (A, rows) and member genes (B, columns) associated (green) with each factor (columns) by their outlier weights in the mixing matrix or outlier loadings in the matrix of source signal estimates, respectively. **C-E.** Choice of number of factors. ‘gn’-main criterion for the ladle estimate ^146^ (C, y-axis, **STAR Methods**), explained variance (**D**, R^2^, y-axis) after matrix reconstruction with estimated components, and distribution of explained variance (**E**, y axis) in randomly sampled unseen perturbation responses (**STAR Methods**) for ICA decomposition with different numbers of components (x-axis). **F.** Factor expression across cells. UMAP embedding of cell profiles (as in Figure 1C) colored by expression scores for each sub-factor (label on top). **G,H.** Jaccard index (color bar) for each pairs of factors (columns and rows) based on genes with outlier loadings in the matrix of source signal estimates (**G**) or regulators with outlier weights in the mixing matrix per factor (**H**).

**Figure S6, related to Figure 6.**
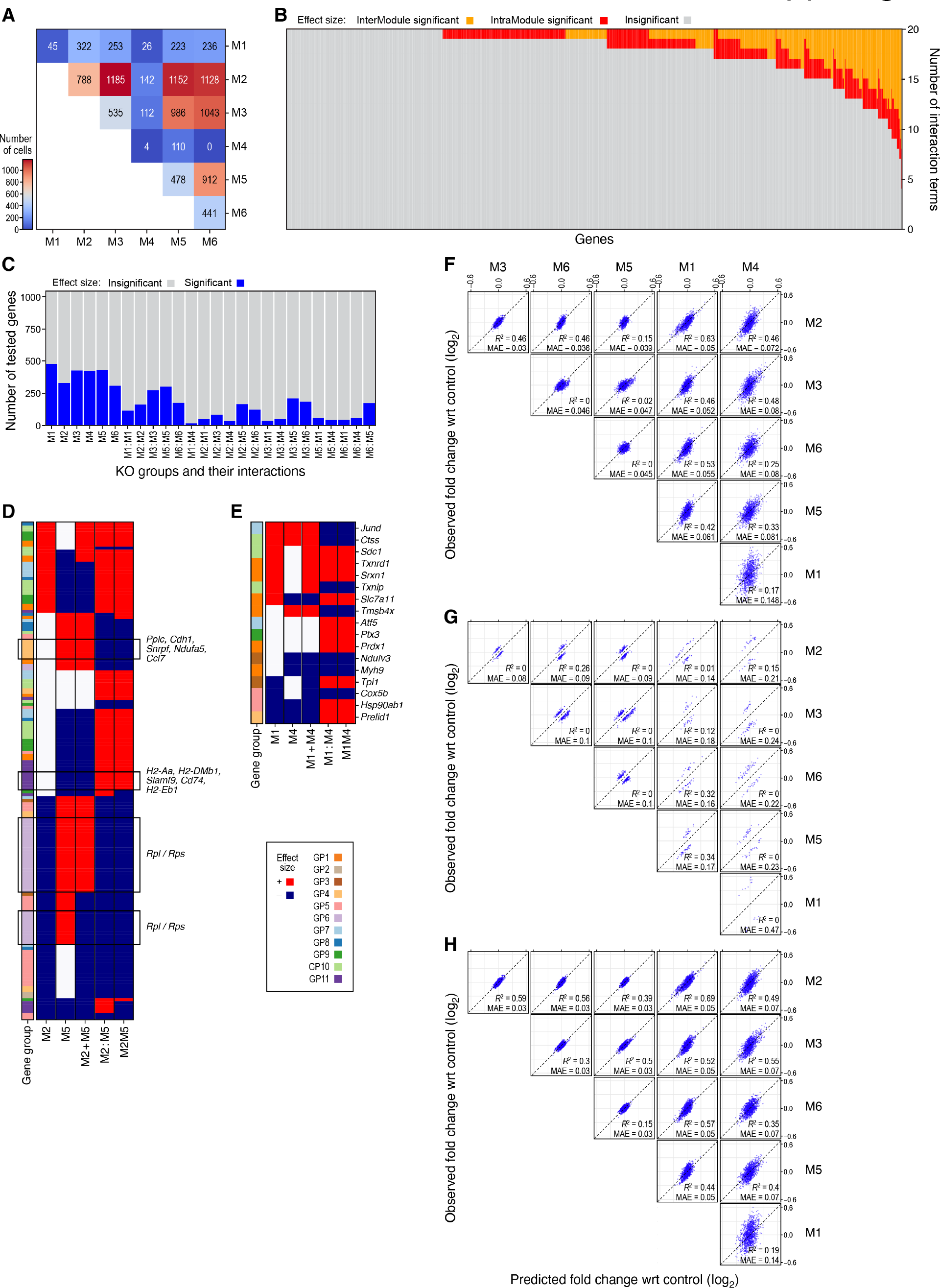
Inter- and intra-module genetic interactions **A.** Combinatorial perturbations in the E3 screen. Number of cells (color bar, number) with pairs of guides targeting genes in the same or pair of modules (rows, columns). **B.** Genes vary in the extent of genetic interactions on their expression. Number of significant interaction terms (FDR < 0.1, y-axis) on the expression of each gene (x axis) from combinatorial perturbations across all intra-modules (red) and inter- (orange) module interactions. **C.** Module pairs vary in the extent of interactions. Number of target genes (y axis, of 1,041 in the regulatory model) with a significant effect on expression (blue) due to single perturbations in genes in one module (M_i_, x axis) or with a significant interaction term due to perturbation in two genes from the same (M_i_:M_i_, x axis) or different (M_i_:M_j_, i≠j, x axis). **D,E.** Specific inter-module interactions across genes. Binarized significant effect (FDR < 0.1) (red/blue: positive/negative) on the expression of genes (row, only genes with significant interaction terms) by single perturbations in regulators from two different modules (M_i_, M_j_, i≠j), their additive effect (M_i_+M_j_), their interaction term (M_i_:M_j_), and the observed effect (columns). Gene program membership is labeled on left. **F-H.** Observed vs. additive effects across genes. Fold changes in gene expression observed (*y* axis) following inter-module combinatorial perturbation or predicted by an additive model (*x* axis) for each of the 1,041 genes (F, dots) or only genes with either significant (G, FDR < 0.1) or non-significant (H, FDR >= 0.1) inter-module interaction terms. R^2^: explained variance in observed fold changes; MAE: mean absolute error of the predictions.

**Figure S7, related to Figure 6.**
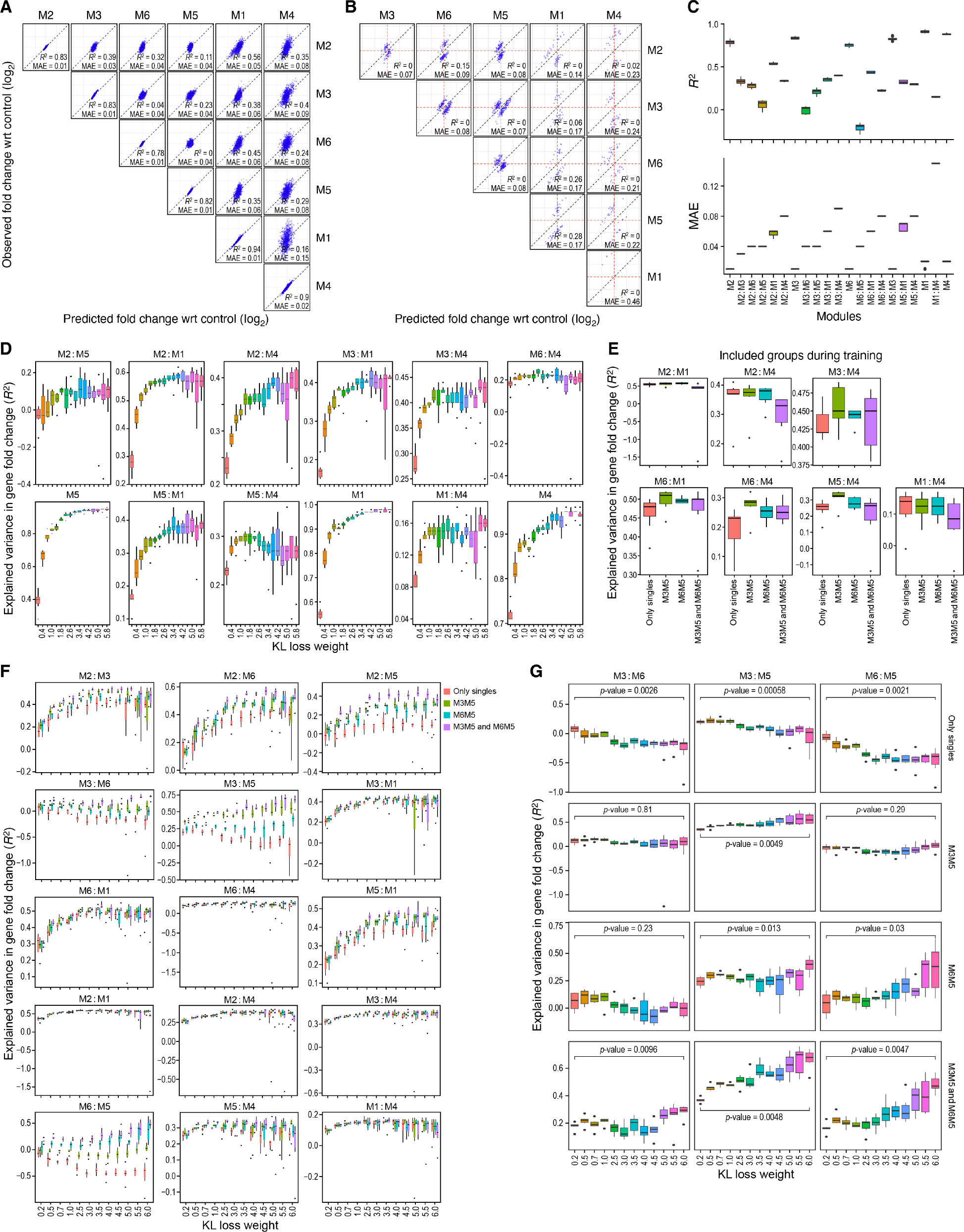
**Com**β**VAE for prediction of combinatorial perturbations A,B.** Fold change predictions from comβVAE. Fold changes in gene expression observed (*y* axis) following inter-module combinatorial perturbation (y-axis) or predicted by comβVAE (*x* axis) for each of the 1,041 genes (A, dots) or only genes with significant (B, FDR < 0.1) inter-module interaction terms. The diagonal entries in (A) reflect the prediction in single knockouts. **C.** comβVAE performance for different modules and module combinations. Distribution of explained variance (top, y axis, R^2^) and mean absolute error (bottom, y axis, MAE) of the predictions of the comβVAE model for each module (M_i_) or inter-module combination (M_i_:M_j_, i≠j) (x axis) across 7 runs with the same hyperparameters. **D.** Impact of changes in KL loss weight. Distribution of explained variance in fold changes (D, y-axis) of the comβVAE at different KL loss weight values (*x* axis) for each module (M_i_) or inter-module combination (M_i_:M_j_, i≠j) across 7 runs with the same hyperparameters. **E-G.** Impact of training with doubly perturbed cells. **E.** Distribution of the explained variance in fold changes (y axis, R^2^) in the indicated inter-module combinations (labels on top of panel) when the model (Beta=6.0) is trained only with data from singly perturbed cells from all modules (M) or singly perturbed cells from all modules and doubly perturbed cells from one or two pairs of modules (M_i_:M_j_; i≠j) (x axis) across 7 runs with the same hyperparameters. **F.** Explained variance in fold changes (y-axis) in each module pair (panels) when the comβVAE is learned with different KL loss weight values (*x* axis) and trained either only with singly perturbed cells (red) or with both singly perturbed and doubly perturbed cells of specific module pairs (green: M3M5, blue: M5M6, purple: M3M5 and M5M6). **G.** Explained variance in fold changes (y-axis) in select module pairs with relatively high number of genes with significant inter-module interaction terms (column headers) when the comβVAE is learned at different KL loss weight values (*x* axis) and ..+ trained either only with singly perturbed cells (red) or with both singly perturbed and doubly perturbed cells of specific module pairs (row labels). All boxes in Box plots display the first (Q_1_), second (Q_2,_median) and third (Q_3_) quartiles, and bottom and top whiskers show the intervals [Q_1_ -1.5 IQR, Q_1_] and [Q_3,_ Q_3_ +1.5 IQR], respectively.

